# Single-cell profiling reveals periventricular CD56^bright^ NK cell accumulation in multiple sclerosis

**DOI:** 10.1101/2021.09.17.460741

**Authors:** Sabela Rodríguez-Lorenzo, Lynn van Olst, Carla Rodriguez-Mogeda, Alwin Kamermans, Susanne M.A. van der Pol, Ernesto Rodríguez, Gijs Kooij, Helga E. de Vries

## Abstract

Multiple sclerosis (MS) is a chronic demyelinating disease characterised by immune cell infiltration resulting in lesions that preferentially affect periventricular areas of the brain. Despite research efforts to define the role of various immune cells in MS pathogenesis, the focus has been on a few immune cell populations while full-spectrum analysis, encompassing others such as natural killer (NK) cells, has not been performed. Here, we used single-cell mass cytometry (CyTOF) to profile the immune landscape of brain periventricular areas – septum and choroid plexus – and blood from MS donors and controls with and without other neurological diseases. Using a 37-marker panel, we revealed the infiltration of T cells and antibody-secreting cells in periventricular brain regions and identified a novel NK cell signature specific to MS. CD56^bright^ NK cells accumulated in the septum of MS donors, displaying an activated and migratory phenotype that was similar to that of CD56^bright^ NK cells in their circulation. We validated this signature by multiplex immunohistochemistry and found that NK cells with high expression of granzyme K, which is typical of the CD56^bright^ subset, accumulated in both periventricular lesions and the choroid plexus. Together, our multi-tissue single-cell data suggests that CD56^bright^ NK cells infiltrate the periventricular brain regions in MS patients via both the blood-brain and blood-CSF barriers and brings NK cells to the spotlight of MS pathology.

## Introduction

Multiple sclerosis (MS) is a chronic neuroinflammatory disease characterised by demyelinating lesions within the central nervous system (CNS). In MS, peripheral immune cells gain access to the CNS and cause severe neuroinflammation, myelin damage, and subsequent neurodegeneration. In the past decades, a wealth of knowledge has been gained on the role of monocyte-derived macrophages, CD8^+^ and CD4^+^ T cells, B cells and antibody-secreting cells in MS pathogenesis [32, 62]. To date, however, we still know relatively little about the presence and roles of other immune cell subsets in the MS brain, such as natural killer (NK) cells.

Brain regions around the ventricles are hotspots for MS lesion formation [8,21,38,50] but the reasons for periventricular susceptibility are poorly understood [40]. Since the majority of periventricular MS lesions occur around a central vessel [1, 55], it has been suggested that vascular topography may influence MS pathology [33]. Although the accumulation of immune cell infiltrates around post-capillary venules suggests their trafficking across the blood-brain barrier [52], periventricular veins drain to the cerebrospinal fluid (CSF) [41]. Thus, MS periventricular pathology may be related to factors from both the blood and the CSF.

In MS, the CSF that flows through the ventricles is enriched in immune cells [9,46,48] and inflammatory factors [24, 61]. The main source of CSF is the choroid plexus, a secretory tissue located in the brain ventricles that acts as an immunological hub [54] and forms the blood-CSF barrier. Thus, the location and functions of the choroid plexus are strategic to regulate periventricular homeostasis and thereby influence neuroinflammation [15, 35]. For example, immune cells infiltrate the brain through the choroid plexus in the early stages in an MS animal model [43]. In MS patients, we and others have shown that immune cells accumulate at the choroid plexus in the progressive phases of the disease [45, 60]. Therefore, we postulate that CSF-mediated immune processes originating in the choroid plexus could participate in the periventricular inflammation typical for MS patients. However, it is still uncertain whether such processes involve cell trafficking into the CNS and/or the secretion of inflammatory factors.

Here, we used single-cell mass cytometry (CyTOF) to profile the immune landscape of periventricular MS brain regions and to better understand their susceptibility and the routes of immune cell infiltration from the periphery into the CNS. For this purpose, we isolated immune cells from the human post-mortem periventricular brain areas (septum), the choroid plexus and the blood. With our 37-marker panel, we defined the main innate and adaptive immune cell populations and their phenotype in MS donors and controls with dementia or without neurological disease. Besides detecting the accumulation of T cells and antibody-secreting cells typical of MS in the periventricular brain regions, we also identified an NK signature consisting of CD56^bright^ NK cells. CD56^bright^ NK cells are known by their immunoregulatory properties and were also detected in the blood of the same MS donors with a migratory and activated phenotype, similarly to those in the brain. Using multispectral immunofluorescence, we validated these findings in an independent cohort of periventricular brain and choroid plexus tissues, which indicated that NK cells expressing granzyme K, typical of the CD56^bright^ subset, are enriched in MS lesions and the choroidal tissue from MS donors. Thus, our data suggests both direct and indirect infiltration across the brain barriers into periventricular areas of the MS brain. Together, our study brings NK cells back to the spotlight of MS pathology by suggesting a local immunoregulatory role of the CD56^bright^ subset within periventricular MS brain regions.

## Results

### Single-cell mass cytometry of the septum reveals an accumulation of T cells and natural killer cells in multiple sclerosis

To investigate the periventricular immune landscape in MS, we performed cytometry by time of flight (CyTOF) on post-mortem septum – a periventricular brain region highly exposed to CSF –, choroid plexus – the main producer of CSF –, and peripheral blood from non-neurological controls, neurological controls diagnosed with dementia and MS donors (Fig. 1a). Demographic and clinical information of the patients is summarised in Table 1 and Supplementary Figs. 1a, 4a and 6a. We used a 37-antibody panel consisting of lineage markers, focused on lymphoid subsets, and phenotypic markers to determine the migratory, activation and memory phenotypes (Table 2). We performed unsupervised clustering for each tissue followed by manual merging of the clusters based on biological knowledge (Supplementary Fig. 1b-e).

**Fig. 1.**
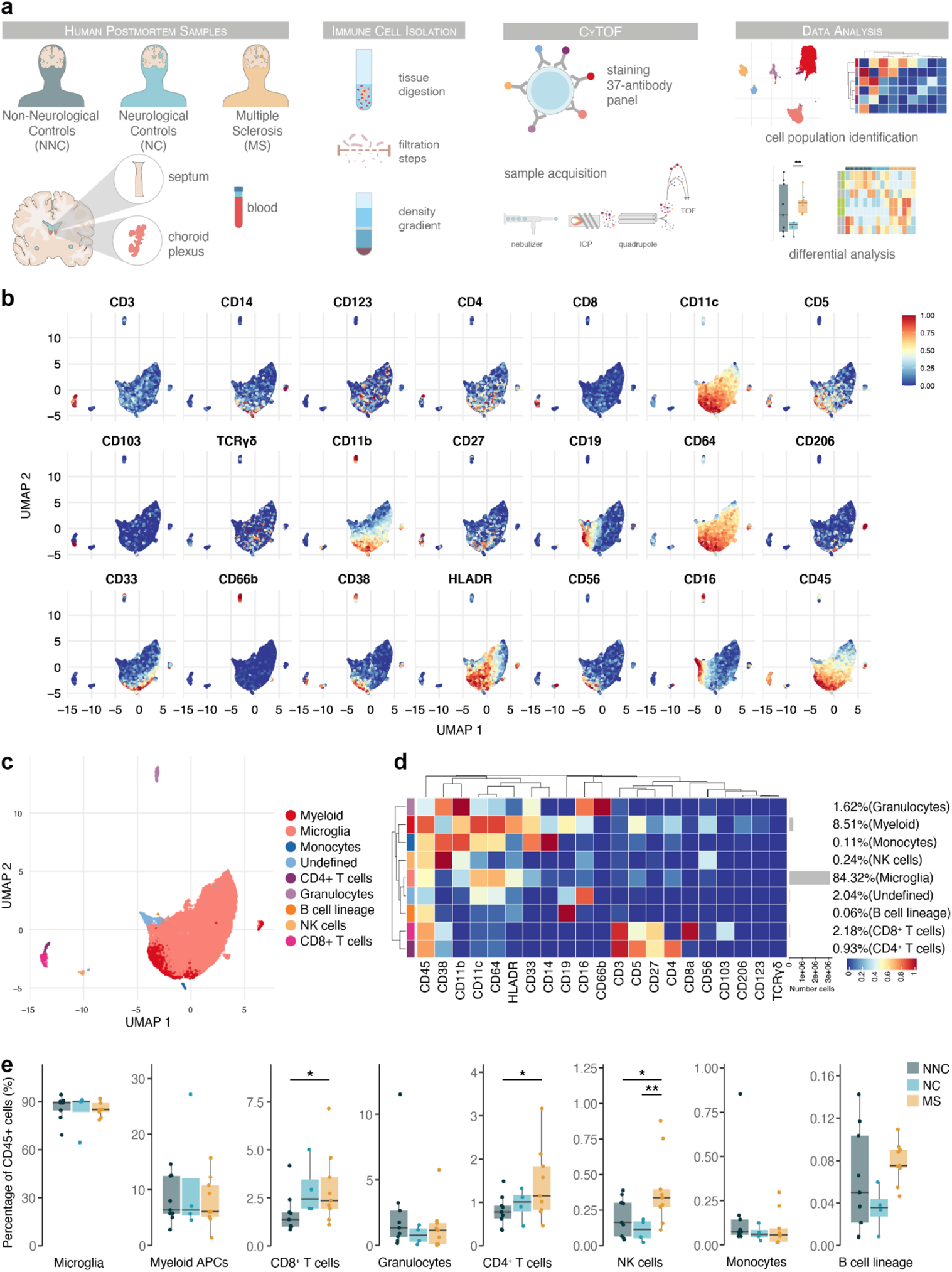
Immune phenotyping of the septum using mass cytometry reveals inflammation in multiple sclerosis involving T cells and natural killer cells. **(a)** Schematic overview of the study design. Immune cells were isolated from septum, choroid plexus and blood samples. Cells were then fixed, barcoded, stained and acquired on a CyTOF Helios system and analysed using FlowSOM and ConsensusClusterPlus. **(b)** UMAP plots based on the arcsinh-transformed expression of the “type” (lineage) markers in the septum-derived immune cells. A subset of 1000 randomly selected cells per sample is shown, coloured according to the expression level of each marker. **(c)** UMAP plot based on the arcsinh-transformed expression of the “type” (lineage) markers in the septum-derived immune cells. A subset of 1000 randomly selected cells is shown per sample, coloured according to the manually annotated clusters. **(d)** Median scaled intensities of the “type” (lineage) markers across the nine annotated septum-derived immune cell populations. The horizontal grey bars show the percentage of each cluster out of the total number of cells. **(e)** Percentage of each annotated cell population out of the total CD45^+^ cells from the septum of non-neurological controls (NNC), neurological controls (NC) and multiple sclerosis (MS) donors. * adjusted P < 0.1; ** adjusted P < 0.05.

Overall, in the septum, we identified 9 immune cell populations (Fig. 1b-d) and one cluster of non-immune cells of possible neuronal origin (CD45^−c^CD56^bright^) (Supplementary Fig. 1e), which was excluded from further analysis. The septum was mainly populated by microglia (84.3 % of CD45^+^ immune cells), followed by other antigen-presenting cells (APCs) of myeloid origin (8.5 %). The frequency of CD8^+^ T cells (2.2 %) more than doubled that of CD4^+^ T cells (0.9 %). Other immune cells present in the septum were granulocytes, natural killer (NK) cells, monocytes, and cells from the B lymphocyte lineage.

We next compared the proportions of each population in the septum among the MS and control groups (Fig. 1e). We found a higher percentage of both CD8^+^ and CD4^+^ T cells in MS compared to non-neurological controls. CD4^+^ T cell accumulation was specific for MS, while the percentage of CD8^+^ T cells was higher in both MS and neurological controls compared to non-neurological controls. We also found a higher percentage of NK cells in the MS group compared to the non-neurological and neurological controls. A moderate negative correlation was found between the percentage of CD8^+^ T cells and age, while there was a moderate positive correlation between the B cell lineage frequencies and postmortem delay (PMD) (Supplementary Fig. 1f).

Thus, we were able to identify the canonical brain immune cell populations in lesion-prone, periventricular brain regions using CyTOF. We found that the septum also presents an accumulation of CD8^+^ and CD4^+^ T cells typical of MS brains [2, 32]. Moreover, we show that NK cell accumulation is a signature of the septum of MS patients.

### Phenotyping the T cell populations in the septum

Next, we investigated which subsets within CD8^+^ and CD4^+^ T cells were accumulating in the septum of MS donors. Briefly, CD8^+^ and CD4^+^ T cells were further subdivided into 10 populations and a tentative biological name was assigned to each of them (Fig. 2a). The septum was mostly populated by CD69^+^ tissue-resident memory T cells (T_RM_), belonging to both CD8^+^ (clusters CD8 c1 and CD8 c2, the latter expressing residency marker CD103) and CD4^+^ T cells (cluster CD4 c1). Effector memory T cells (T_EM_) were also present (clusters CD8 c4 and CD4 c2), including a cluster of CD8^+^ T cells re-expressing CD45RA (T_EMRA_ cells, cluster CD8 c3). A small cluster of CD8^+^ T cells presented an intermediate or transitional phenotype between naïve and memory (cluster CD8 c5). Naïve CD4^+^ T cells (cluster CD4 c3) and CD4^+^ T_REGS_ (cluster CD4 c4) were present in small proportions. We also identified a small cluster of γδ T cells, which was not detected in the general clustering of Fig. 1.

**Fig. 2.**
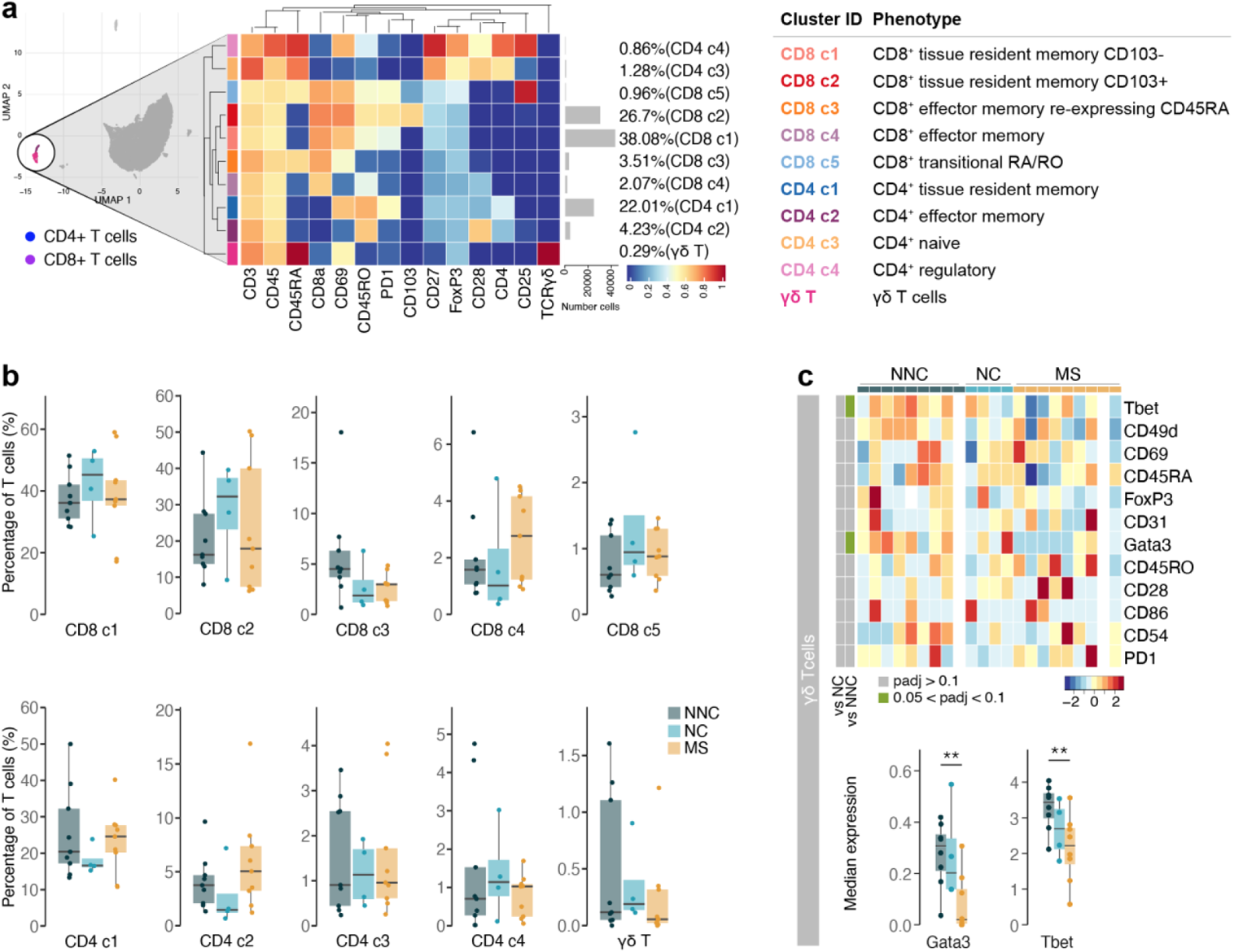
Phenotyping the T cell populations in the septum using mass cytometry. **(a)** Median scaled intensities of the “type” markers across T cell subpopulations in the septum. The horizontal grey bars show the percentage of each cluster out of the total investigated cells. The table on the right shows the tentative biological names of the T cell subpopulations. **(b)** Percentage of each annotated cell population out of the total number of T cells (CD4^+^ and CD8^+^ T cells) from the septum of NNC, NC and MS donors. **(c)** Median scaled intensities of the “state” markers in γδ T cells across all septum samples. Column annotation shows disease groups; row annotation shows adjusted p-values of comparing MS vs NC and MS vs NNC samples. Colour key shows row Z score. Blank cells are shown in a sample where γδ T cells were absent. Boxplots show median expression in γδ T cells of selected markers showing differential expression between the MS and NNC samples. ** adjusted P < 0.05. NNC: non-neurological controls; NC: neurological controls; MS: multiple sclerosis.

Proportions of the different T cell subsets present in the septum did not differ among the disease groups (Fig. 2b). However, before correcting for multiple testing, the percentage of CD4^+^ T_EM_ cells was higher in MS relative to neurological controls (unadjusted P = 0.03), while the proportion of CD8^+^ T_EMRA_ cells was lower in both disease groups than in non-neurological controls (MS vs. non-neurological controls: unadjusted P = 0.04; neurological controls vs. non-neurological controls: unadjusted P = 0.09). Interestingly, γδ T cells in MS patients displayed reduced expression of the transcription factors T-bet and GATA3 compared to the controls (Fig. 2c), suggesting a less activated phenotype [65].

Thus, similarly to other brain areas [53], the septum is mainly comprised of tissue-resident memory cells and, to a lesser extent, effector memory T cells. The proportions of T cells subsets were equal between MS and controls, suggesting that the accumulation of T cells in MS (Fig. 1e) does not result from the enrichment of a particular T cell subset we were able to detect but from an accumulation of all of them.

### Accumulation of activated CD56^bright^ NK cells and antibody-secreting cells in the MS septum

The higher percentage of NK cells observed in MS led us to further investigate this cell population using markers for cell migration and activation. Phenotypically, NK cells from MS patients displayed a more migratory profile than those from non-neurological and neurological controls (Fig. 3a): higher expression of CD49d (integrin alpha-4), CD54 (ICAM1 or intercellular adhesion molecule 1) and CD31 (PECAM1 or platelet endothelial cell adhesion molecule). Contrarily, the expression of CD45RA and T-bet, markers of immature and cytotoxic NK cells respectively [6, 56], was lower in NK cells from MS donors compared to both control groups.

**Fig. 3.**
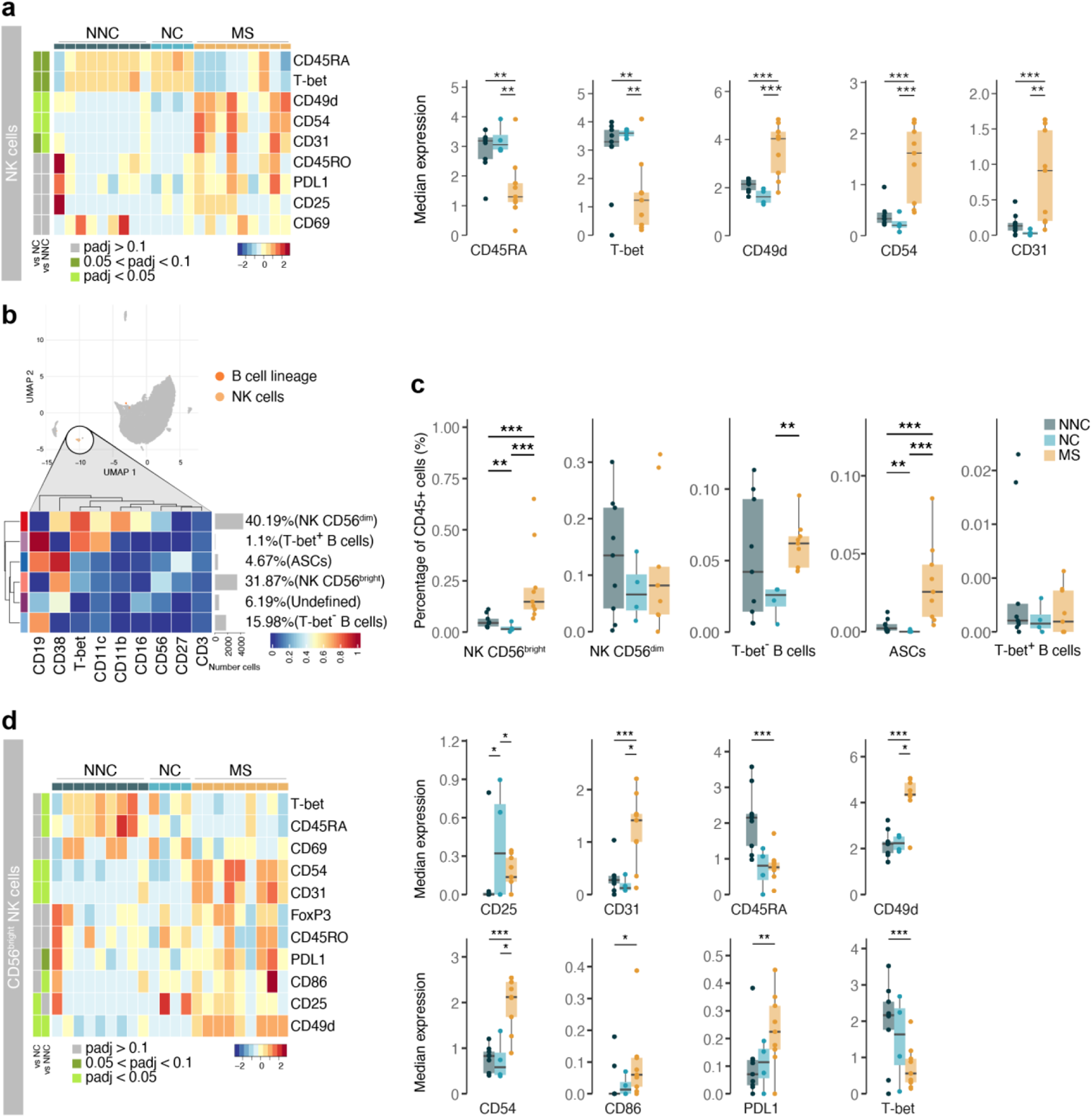
Activated CD56^bright^ NK cells and antibody-secreting cells accumulate in the MS septum. **(a)** Median scaled intensities of the “state” markers in NK cells across all septum samples. Column annotation shows disease groups; row annotation shows adjusted p-values of comparing MS vs NC and MS vs NNC. Colour key shows row Z score. Boxplots show median expression in NK cells of markers with differential expression between MS and NNC samples. **(b)** Median scaled intensities of the “type” markers across NK cell and B cell lineage populations in the septum. The horizontal grey bars show the percentage of each cluster out of the total investigated cells. **(c)** Percentage of each annotated cell population out of the total CD45^+^ cells from the septum of NNC, NC and MS donors. **(d)** Median scaled intensities of the “state” (phenotype) markers in CD56^bright^ NK cells across all septum samples. Column annotation shows disease groups; row annotation shows adjusted p-values of comparing MS vs NC and MS vs NNC samples. Colour key shows row Z score. Boxplots on the right show median expression in CD56^bright^ NK cells of selected markers showing differential expression between the MS and NNC samples. NK: natural killer; ASCs: antibody-secreting cells. * adjusted P < 0.1; ** adjusted P < 0.05, *** adjusted P < 0.01. NNC: non-neurological controls; NC: neurological controls; MS: multiple sclerosis

Next, we further analysed NK cells together with B cells, as they share the expression of markers such as CD38, T-bet and CD27, and divided the two populations into tinier subsets. Two NK cell populations have been described in humans based on their expression of CD56 and CD16 [34]. In the septum, we identified both CD56^dim^ CD16^+^ NK cells and CD56^bright^ CD16^-^ NK cells in similar frequencies, two subsets of B cells (CD19^+^ CD27^-^ CD38^-^ CD11c^-^ T-bet^-^ and CD19^+^ CD27^-^ CD38^-^ CD11c^+^ T-bet^+^) and another of antibody-secreting cells (ASCs; CD19^+^ CD27^+^ CD38^+^) (Fig. 3b and Supplementary Fig. 3a-c).

CD56^bright^ NK cells in the septum were more abundant in MS compared to controls (Fig. 3c). Moreover, CD56^bright^ NK cells in MS donors expressed higher levels of adhesion molecules (CD49d, CD54 and CD31), ligand PDL1, costimulatory molecule CD86, and lower levels of CD45RA and T-bet relative to controls (Fig. 3d). Overall, this is a very similar profile to that observed in the general NK cell cluster (Fig. 3a), suggesting that accumulation of CD56^bright^ NK cells in MS septum led to the observed increase in NK cells in the general clustering of septum-derived immune cells.

We also observed changes in the B cell lineage of the MS septum (Fig. 3c). The frequency of B cells was higher in MS relative to neurological controls, but not relative to non-neurological controls. Of note, B cells in neurological controls (with dementia) displayed an activated phenotype with higher expression of CD25 and CD49d (Supplementary Fig. 3d). ASCs were rare in both control groups and were a unique feature of the MS septum (Fig. 3c).

Together, our data suggests that in MS, CD56^bright^ NK cells either infiltrate the brain more efficiently than their CD56^dim^ counterpart and/or proliferate at the septum leading to CD56^bright^ NK cell accumulation. Moreover, we show that ASCs are also a typical feature of the MS septum, as seen in other brain regions [32].

### Activated CD56^dim^ NK cells at the choroid plexus in MS

The central location of the septum adjacent to the brain ventricles guarantees a high exposure to the CSF. In MS patients, the immune environment in the CSF is altered [9, 46]. Here, we wanted to investigate if immune cell alterations in the MS septum correlate with those in the main producer of CSF: the choroid plexus.

First, we wanted to identify and characterize the main immune cell populations in the choroid plexus. Overall, we identified 9 immune cell populations (Fig. 4a-b), with myeloid APCs (macrophages and dendritic cells) being the most abundant, followed by granulocytes. We also detected CD8^+^ and CD4^+^ T cells, NK cells, monocytes and cells from the B cell lineage. Unlike at the septum, the frequencies of the main immune cell subsets at the choroid plexus were similar among the disease groups (Fig. 4c). We found the choroid plexus to be populated with a variety of T cells, the most abundant subset being CD8^+^ tissue-resident memory T cells (CD8 c1) (Fig. 5a and Supplementary Fig. 5a). Unlike the subset of CD8^+^ CD103^+^ T_RM_ cells from the septum (cluster CD8 c2), those in the choroid plexus did not express CD103 (cluster CD8 c1). We also identified double-negative T cells, but we could not detect CD4^+^ T_REGS,_ suggesting a very low abundance of these cells in the choroid plexus. As seen in the septum, all disease groups showed a similar frequency of the T cell subsets (Supplementary Fig. 5b).

**Fig. 4.**
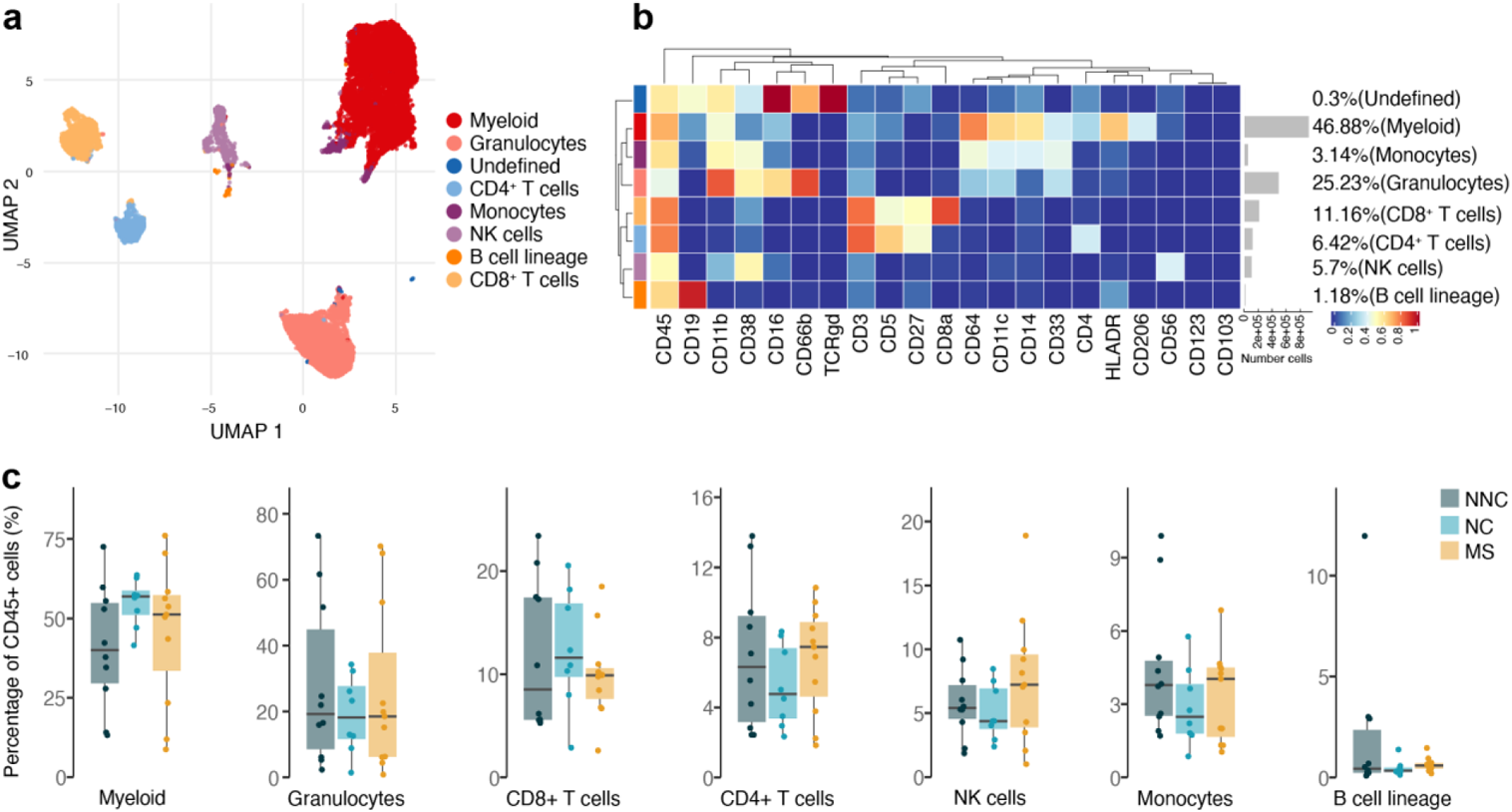
Immune phenotyping of the choroid plexus using mass cytometry suggests NK cell infiltration in MS. **(a)** UMAP plot based on the arcsinh-transformed expression of the “type” (lineage) markers in the choroid plexus-derived immune cells. A subset of 1000 randomly selected cells per sample is shown, coloured according to the manually annotated clusters. **(b)** Median scaled intensities of the “type” (lineage) markers across choroid plexus-derived immune cell populations. The horizontal grey bars show the percentage of each cluster out of the total number of cells. **(c)** Percentage of each annotated cell population out of the total number of CD45^+^ cells from the choroid plexus of NNC, NC and MS donors. NNC: non-neurological controls; NC: neurological controls; MS: multiple sclerosis

**Fig. 5.**
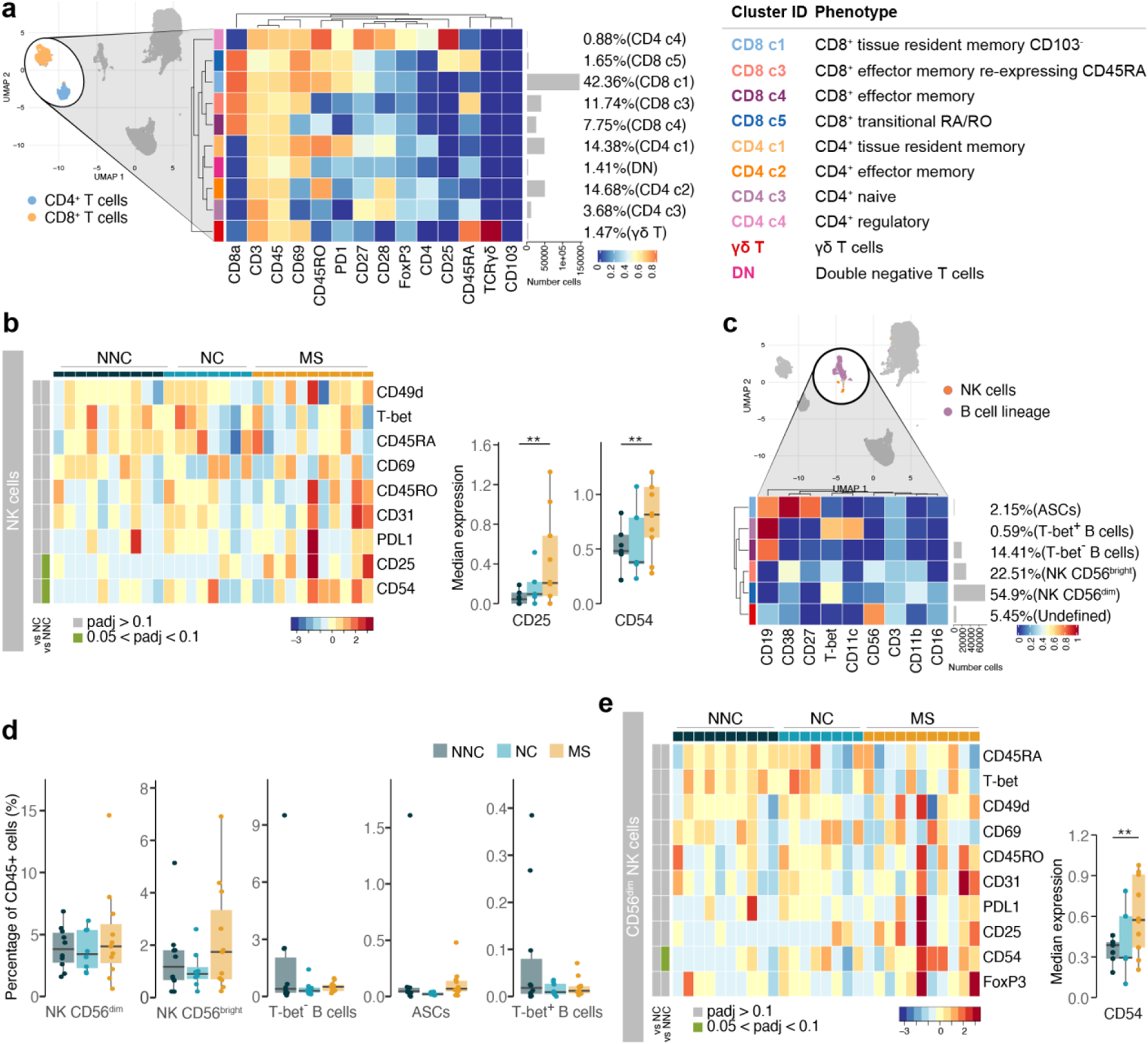
The choroid plexus in MS presents alterations in the NK cell populations. **(a)** Median scaled intensities of the “type” markers across T cell subsets in the choroid plexus. The table shows tentative biological names of the T cell subpopulations. **(b)** Median scaled intensities of the “state” markers in NK cells across all choroid plexus samples. Column annotation shows disease groups; row annotation shows adjusted p-values of comparing NC vs NNC and MS vs NNC samples. Colour key shows row Z score. Boxplots show median expression in NK cells of markers with differential expression between MS and NNC. **(c)** Median scaled intensities of the “type” markers across NK cell and B cell lineage populations in the choroid plexus. **(d)** Percentage of each annotated NK or B cell population out of the total number of CD45^+^ cells from the choroid plexus of NNC, NC and MS donors. **(e)** Median scaled intensities of the “state” markers in CD56^dim^ NK cells across all choroid plexus samples. Column annotation shows disease groups; row annotation shows adjusted p-values of comparing NC vs NNC and MS vs NNC samples. Colour key shows row Z score. Boxplot shows median expression in CD56^dim^ NK cells of the marker with differential expression between MS and NNC. ** adjusted P < 0.05. NNC: non-neurological controls; NC: neurological controls; MS: multiple sclerosis **(a, c)** The horizontal grey bars show the percentage out of the total cells.

NK cells from the MS choroid plexus expressed higher levels of the IL-2 receptor α-chain CD25 and of adhesion molecule CD54 relative to those from controls (Fig. 5b). A closer look at the NK and B cell lineages (Fig. 5c and Supplementary Fig. 5c) revealed that CD56^dim^ NK cells were more abundant than CD56^bright^ NK cells in the choroid plexus in all groups. Unlike in the septum, ASCs were rare in the choroid plexus regardless of disease. Although the frequencies of NK or B cell lineage cells were similar among the groups, there was a trend to a higher percentage of CD56^bright^ NK cells in MS donors relative to both control groups (Fig. 5d). However, it was the CD56^dim^ subset that expressed higher levels of CD54 in MS than in controls (Fig. 5e).

In sum, myeloid APCs are the most abundant immune cell population in the choroid plexus. Our high-dimensional approach allowed us to better define the subsets of T cells and other less abundant immune cells such as NK cells. Interestingly, at the MS choroid plexus, we found a phenotype shift in the CD56^dim^ NK cells which did not result in a different abundance in the tissue. While there only was a non-significant trend for more CD56^bright^ NK cells, we cannot discard the possibility that the choroid plexus contributes to the infiltration of CD56^bright^ NK cells in the MS septum.

### Circulating CD56^bright^ NK cells show a migratory and activated phenotype in MS

The high percentage of granulocytes and monocytes detected in the choroid plexus suggested a strong contribution from peripheral cells circulating in its abundant vascular network. Indeed, when immune cells from all tissues were plotted in a UMAP, high similarities between choroid plexus and blood-derived immune cell populations were revealed (Fig. 6a and Supplementary Fig. 6d). The septum, on the contrary, contained mostly brain-resident microglia and other tissue-resident immune cells.

**Fig. 6.**
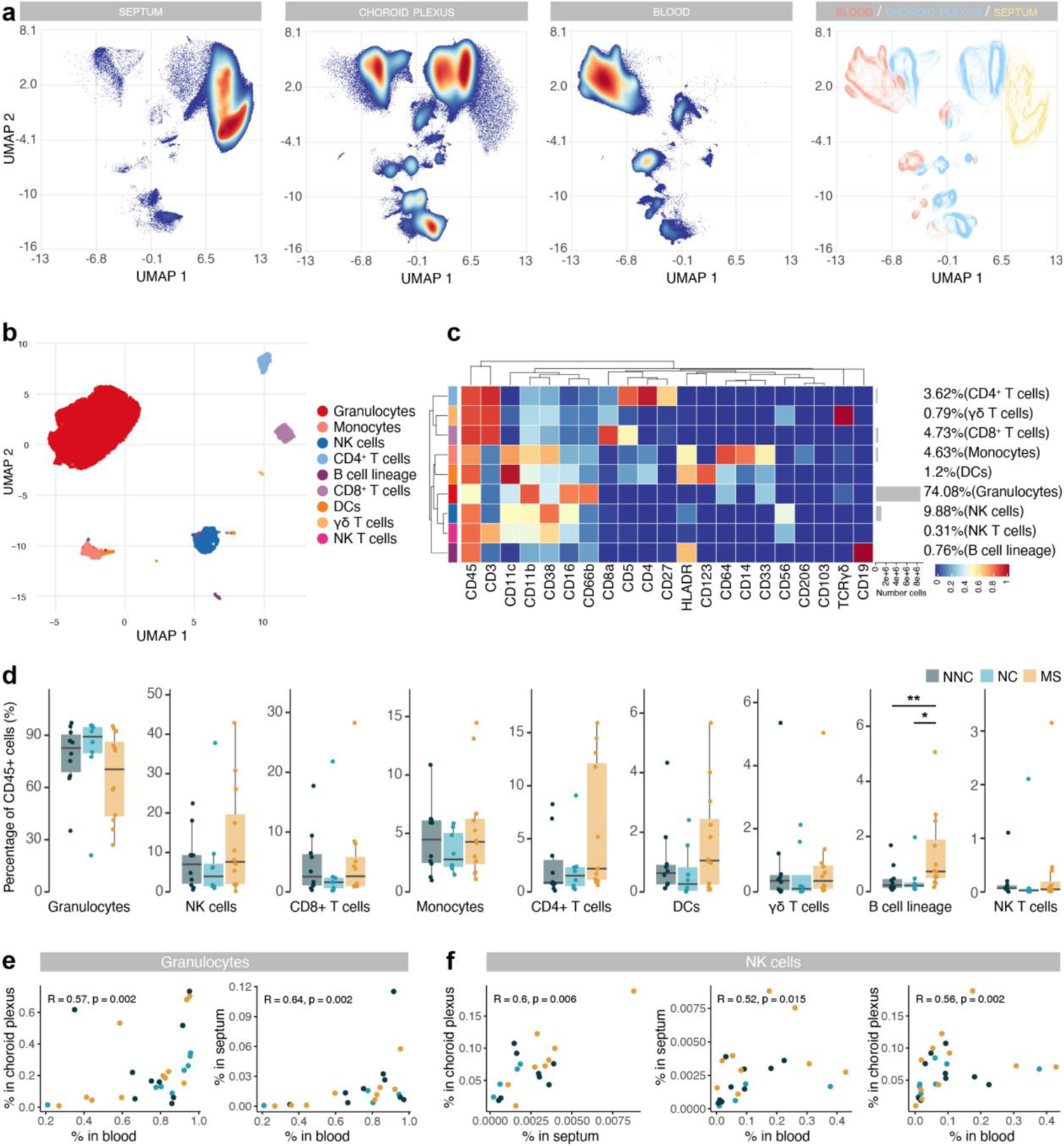
Immune phenotyping reveals an accumulation of B cells in post-mortem MS blood. **(a)** UMAP plot based on the arcsinh-transformed expression of all markers in the immune cells of the septum, choroid plexus and blood. 100,000 randomly selected cells were selected per sample. The first three plots are coloured according to the density. The final plot shows an overlay figure where cells are coloured by their tissue of origin. **(b)** UMAP plot based on the arcsinh-transformed expression of the “type” (lineage) markers in the blood-derived immune cells. A subset of 1,000 randomly selected cells per sample is shown, coloured according to the manually annotated clusters. **(c)** Median scaled intensities of the “type” markers across blood-derived immune cell populations. Horizontal grey bars show the percentage out of the total cells. (**d)** Percentage of each annotated cell population out of the total number of CD45^+^ cells from the blood of NNC, NC and MS donors. **(e)** Scatter plots showing correlations between the proportions of granulocytes in blood, choroid plexus and septum. Each dot represents a sample, coloured by disease group. **(f)** Scatter plots showing correlations between the proportions of NK cells in different tissues. Each dot represents a sample, coloured by disease group. R: Spearman’s Rho rank correlation coefficient; p = p-value. * adjusted P < 0.1; ** adjusted P < 0.05. NNC: non-neurological controls; NC: neurological controls; MS: multiple sclerosis.

In blood, granulocytes comprised the main cell type (Fig. 6b-d and Supplementary Fig. 6e). NK cells were the second most abundant population, followed by T cells and monocytes. Of note, T cell subsets were present at a CD4:CD8 ratio of 0.76:1, instead of the expected 2:1, possibly due to the post-mortem delay [12]. Circulating B cells were more frequent in MS blood compared to controls (Fig. 6d). The frequency of NK cells in the choroid plexus positively correlated with that in the septum and blood (Fig. 6f and Supplementary Fig. 6f), opening up the possibility that NK cells traffic through the blood-CSF barrier.

We detected different populations of circulating T cells, including some that were absent in tissues such as CD8^dim^ T cells (cluster CD8 c6), central memory CD4^+^ T cells (T_CM_, cluster CD4 c5) and CD4^+^ T_EMRA_ (cluster CD4 c6) (Fig. 7a and Supplementary Fig. 7a-b). The shift in B cells in MS was mostly due to T-bet^-^ B cells (Fig. 7b-c), but the same trend was observed in T-bet^+^ B cells (MS vs. non-neurological controls: unadjusted P = 0.06; MS vs. non-neurological controls: unadjusted P = 0.09). Interestingly, T-bet^+^ B cells expressed higher levels of T-bet and CD31 in MS relative to controls (Fig. 7d).

**Fig. 7.**
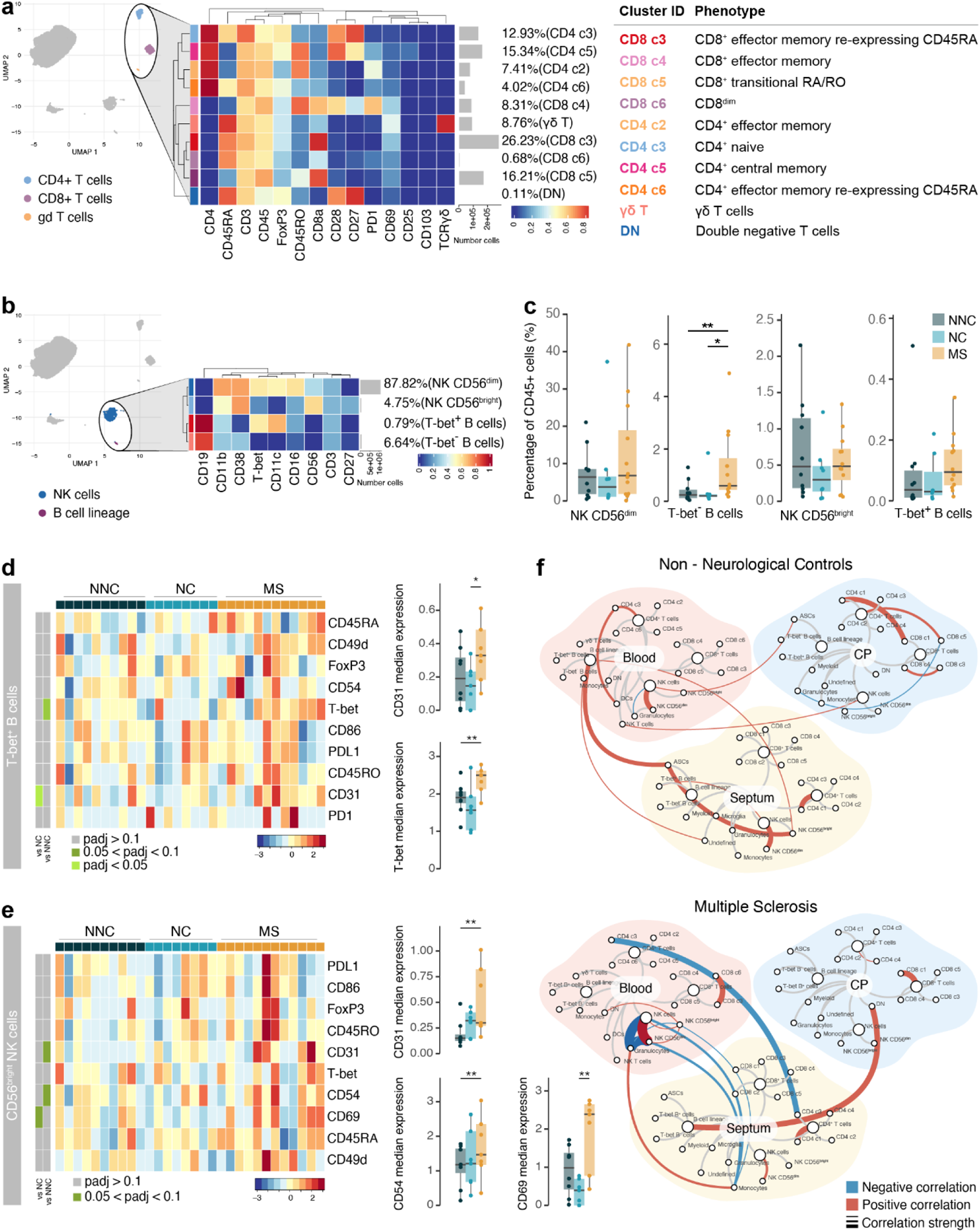
Circulating Tbet^+^ B cells in MS express higher levels of T-bet. **(a)** Median scaled intensities of the “type” markers across T cell subpopulations in blood. Table shows tentative biological names of the T cell subpopulations. **(b)** Median scaled intensities of the “type” markers across NK cell and B cell lineage populations in blood. **(c)** Percentage of NK and B cell lineage populations out of the total CD45^+^ cells from the blood of NNC, NC and MS donors. **(d)** Median scaled intensities of the “state” markers in T-bet^+^ B cells across all blood samples. Column annotation shows disease groups; row annotation shows adjusted p-values of comparing NC vs NNC and MS vs NNC. Colour key shows row Z score. Boxplots show median expression in T-bet^+^ B cells of markers with differential expression between MS and NNC. **(e)** Median scaled intensities of “state” markers in CD56^bright^ NK cells across all blood samples. Column annotation shows disease groups; row annotation shows adjusted p-values of comparing NC vs NNC and MS vs NNC samples. Colour key shows row Z score. Boxplots show median expression in CD56^bright^ NK cells of markers with differential expression between MS and NNC. **(f)** Network plot illustrating correlations between identified immune subsets in the three anatomical sites analysed. Positive correlations are displayed in red and negative correlations in blue. Width of the line is proportional to the correlation coefficients. P-values adjusted with Benjamini-Hochberg method. ** adjusted P < 0.05. NNC: non-neurological controls; NC: neurological controls; MS: multiple sclerosis **(a, b)** Horizontal grey bars show the percentage out of the total cells.

Most NK cells in blood were CD56^dim^ (Fig. 7b-c and Supplementary Fig. 7c), and their percentage strongly correlated with that from the choroid plexus (Supplementary Fig. 7d). The abundance of NK cells was similar among disease groups, but the CD56^bright^ NK cluster from MS blood expressed higher levels of the adhesion molecules CD54 and CD31 (Fig. 7e), in line with the phenotype of CD56^bright^ NK cells in MS septum. This may indicate preferential migration of the CD56^bright^ subset from blood to the septum.

Correlations of immune cell abundance between the different tissues revealed a different pattern in non-neurological controls than in MS (Fig. 7f). In MS donors, there were more connections between immune cells derived from the septum, choroid plexus and the blood, suggesting increased cellular communication and movement in periventricular areas in MS. For example, naïve CD4 T cells (CD4 c3) from blood negatively associated with naïve CD4 T cells in the MS septum but not in non-neurological controls, suggesting some form of trade-off mechanism or migration that is not seen in controls.

Overall, despite the high vascularization of the choroid plexus, we observed a different immune profile in the blood compared to that in the choroid plexus. First, the higher frequency of circulating B cells in MS relative to controls was not observed in the choroid plexus; second, the migratory phenotype of CD56^dim^ NK cells was altered in MS choroid plexus but not in MS blood; third, we show that circulating CD56^bright^ NK cells but not their CD56^dim^ counterpart, display a migratory phenotype in MS. In summary, we identified several associations among the immune populations of the septum, the choroid plexus and the blood, indicative of a dynamic immune environment in periventricular tissues.

### Granzyme K^+^ NK cells accumulate in periventricular lesions and choroid plexus from MS donors

Finally, to confirm that CD56^bright^ NK cells accumulate in periventricular brain regions in MS donors, we used multiplex immunohistochemistry in an independent cohort (Table 1). This technique allowed us to stain up to seven markers in the same section. Due to the difficulty of making a clear distinction between bright and dim expression of CD56 in NK cells by immunohistochemistry, we used granzymes (Gr) expression as a surrogate marker. We defined CD56^bright^ NK cells as CD45^+^ NKp46^+^ GrK^+^ GrB^-^ and CD56^dim^ NK cells as CD45^+^ NKp46^+^ GrK^-^ GrB^+^ [7, 18].

We analysed the presence of NK cells in periventricular brain tissue from controls and within normal-appearing white matter (NAWM) and white matter lesions from MS donors. Lesions were identified by the absence of myelin proteolipid protein and the abundance of HLA-DR^+^ cells (Fig. 8a). Both in the border and the lesion centre we observed a higher density of GrK^+^ NK cells compared to NAWM (Fig. 8b-c). Matching the CyTOF data, GrB^+^ NK cells were more abundant than GrK^+^ cells.

**Fig. 8.**
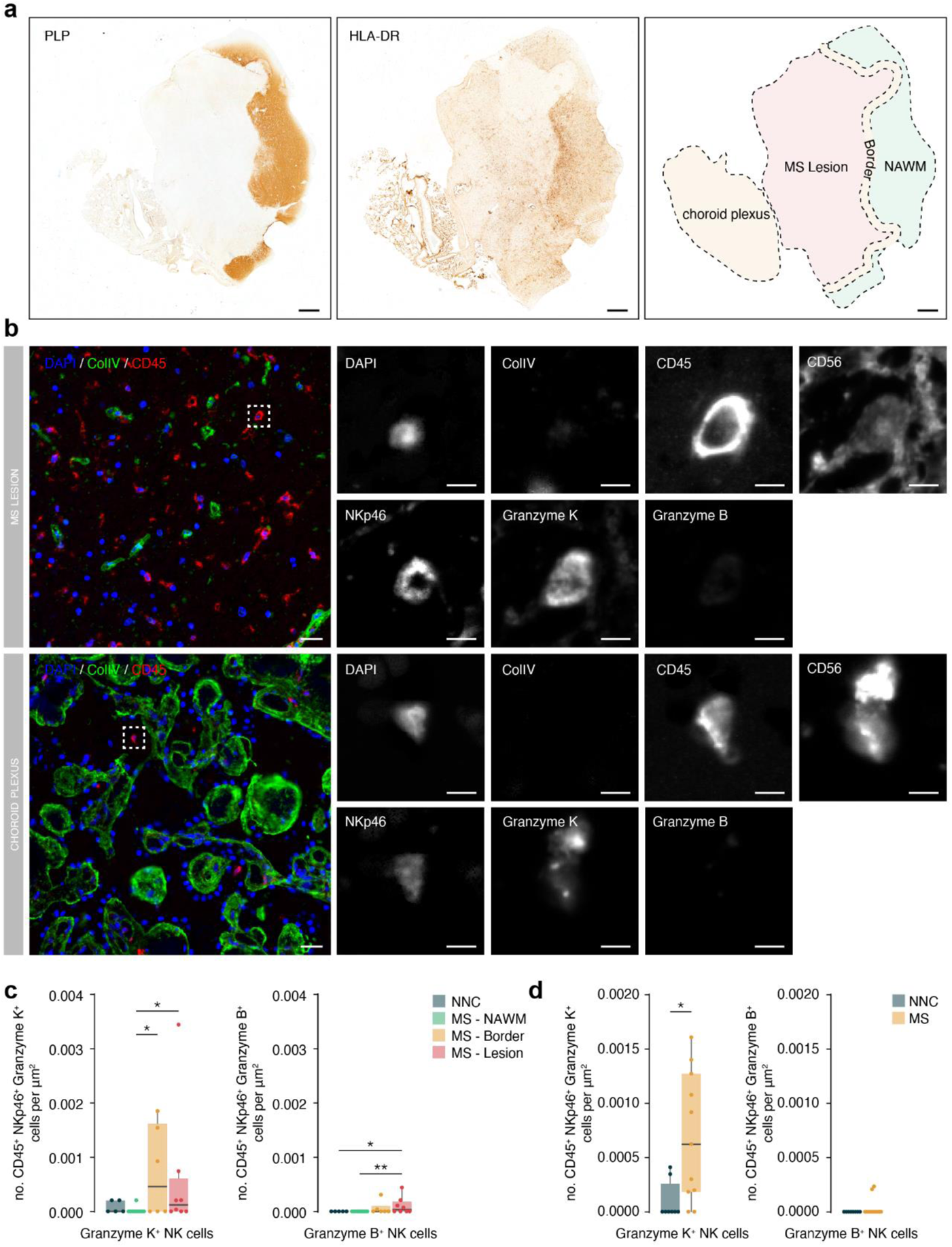
Granzyme K high NK cells accumulate in periventricular lesions and choroid plexus. **(a)** Representative immunohistochemical staining of myelin with proteolipid protein (PLP) and presence of immune cells with HLA-DR. The overview shows the location of the normal-appearing white matter (NAWM), the lesion border, the lesion and the choroid plexus. Scalebar = 1mm. **(b)** Representative multiplex immunohistochemical staining of MS lesion and the choroid plexus. The merged overview shows DAPI (blue), ColIV (green) and CD45 (red). The dotted box indicates CD45^+^ NKp46^+^ Granzyme K^+^ cell. A magnified image of all separate channels is shown in greyscale. Scalebar = 25um for overview and 5um for magnifications. **(c)** Boxplots display the abundance of CD45^+^ NKp46^+^ Granzyme K^+^ and CD45^+^ NKp46^+^ Granzyme B^+^ cells per µm^2^ in MS lesions. NNC: non-neurological controls; MS: multiple sclerosis; NAWM: normal-appearing white matter; * P < 0.05. **(d)** Boxplots display the abundance of CD45^+^ NKp46^+^ Granzyme K^+^ and CD45^+^ NKp46^+^ Granzyme B^+^ cells per µm^2^ in choroid plexus. NNC: non-neurological controls; MS: multiple sclerosis. * P < 0.05.

In addition, the use of choroid plexus tissue sections allowed us to specifically look at infiltrated cells and discard those in the circulation that confounded our CyTOF analysis. In this cohort, we observed a significantly higher density of GrK^+^ but not GrB^+^ NK cells in the choroid plexus of MS vs. control donors (Fig. 8b and d), indicating that the increase shown using CyTOF was specific to the tissue and not due to the high vascularization of the choroid plexus.

In sum, we validated the accumulation of NK cells within the periventricular brain of MS donors. In particular, GrK^+^ NK cells were more abundant and enriched in MS lesions and the choroid plexus stroma.

## Discussion

In this study, we used a CyTOF multi-parameter approach to provide a comprehensive overview of the immune landscape in human periventricular brain tissue from MS and control donors.

Here, we revealed the involvement of CD56^bright^ NK cells in MS periventricular pathology. Since periventricular areas of the brain are a predilection site for MS lesions [8,21,38], we focused on the septum as it separates the two lateral ventricles and is highly exposed to CSF. Although magnetic resonance imaging studies demonstrated the presence of MS lesions in the septum [14,38,39,50], the composition of the inflammatory infiltrates remained unknown. We found an expansion of CD56^bright^ NK cells with an activated and migratory phenotype within the septum of MS donors. Using multiplex immunohistochemistry, we validated the accumulation of NK cells expressing GrK, typical of the CD56^bright^ subset [7], in periventricular MS lesions from an independent cohort. We also found a slight increase in the density of GrB^+^ NK cells — the putative CD56^dim^ cells — in MS lesions versus NAWM and healthy controls that was not detected with the CyTOF. Next to NK cells, we detected higher frequencies of ASCs, CD4^+^ and CD8^+^ T cells in the septum from MS donors compared to controls, which is similar to other periventricular regions [32]. The distribution of general immune cell populations in the septum was consistent with previous literature [26, 53].

NK cells are the lymphocytes of the innate immune system with the ability to regulate adaptive responses. There are two main NK cell subsets based on the expression of CD56 and CD16: CD56^bright^ NK cells express low levels of CD16 and are specialised immunoregulators, while CD56^dim^ NK cells have a potent cytolytic capacity, which may be mediated by CD16 [34]. Importantly, we relied on the markers CD56 and NKp46 to identify human NK cells by CyTOF and immunohistochemistry, respectively. However, NK cells belong to the broader group of innate lymphoid cells (ILCs), which also include ILC1, ILC2 and ILC3 subsets. ILC subpopulations are highly plastic and their identification remains a challenge; it has been proposed that they belong to a single, highly plastic population, but this debate is beyond the scope of this study. Although NK cells are thought to predominate in the brain parenchyma [49], we cannot exclude the presence of other ILCs. Particularly, certain subsets of ILC3s share the expression of CD56 with NK cells [57]. The extent and pathological consequences of the presence of ILC3s in MS should be determined in future studies.

The beneficial role of CD56^bright^ NK cells in MS was first discovered with the anti-inflammatory drug daclizumab. Daclizumab is an anti-CD25 antibody therapy that leads to the expansion of circulating CD56^bright^ NK cells [4, 63], an increase that was later reported with other disease-modifying treatments for MS as well [16,47,51]. In line with this, a protective role for NK cells was shown in animal models of MS [19, 64] through cytotoxicity against autoreactive T cells [64]. Despite the beneficial immunomodulatory role assigned to CD56^bright^ NK cells in the context of MS, the presence of NK cells in the CNS can also be detrimental [23,27,29,30,37]. Still, any potential cytotoxicity of NK cells in the CNS might be halted by tolerogenic signals coming from brain resident cells, such as microglia [31]. Thus, while NK cells may act as a double-edged sword in MS [17], we postulate that the presence of specifically CD56^bright^ NK cells in the periventricular areas primarily has a beneficial effect on MS pathology by limiting neuroinflammation.

To date, studies of NK cells in the MS brain parenchyma did not distinguish between the CD56^bright^ and CD56^dim^ NK subsets. NK cells were detected in the brains from MS donors [17,27,29,58], while absent from controls [17, 29]. Within active MS lesions, NK cells express GrK polarised towards neighbouring T cells [17, 22]. Thus, the expansion of the CD56^bright^ NK cell population we observed in MS brains may mediate a protective response to neuro-inflammation by killing autoreactive T cells through the release of GrK. Moreover, we observed higher expression of the ligand PDL1 in CD56^bright^ NK cells from MS brains compared to those from non-neurological controls, suggesting an involvement of the PD1-PDL1 axis. PDL1 is an NK cell activation marker associated with enhanced cytotoxicity [11] and binds the T cell co-inhibitory molecule PD1 inducing suppression of PD1^+^ autoreactive T cells [44].

Proportions of circulating CD56^bright^ NK cells remained the same in MS and control donors, as described before [17, 28]. Interestingly, CD56^bright^ NK cells in both the MS brain and blood expressed higher levels of proteins associated with cell migration. As such, the enrichment of CD56^bright^ NK cells in the CNS of MS patients might result from selective infiltration from the blood towards the brain rather than being the result of accumulation in the circulation. In line with this, migration studies showed that CD56^bright^ NK cells have a higher capacity to transmigrate across a blood-brain barrier *in vitro* model than their CD56^dim^ counterparts [17]. However, this study did not show enhanced migration of CD56^bright^ NK cells from MS donors compared to controls.

Direct infiltration from the circulation through the local brain vasculature is a likely origin of MS immune infiltrates. Besides, abundant periventricular veins drain to the ventricles [41], resulting in an indirect peripheral-brain connection through the choroid plexus via the CSF. Here, we describe the presence of NK cells in the human choroid plexus and identify a migratory phenotype in CD56^dim^ NK cells in MS. The higher ratio of CD56^bright^/CD56^dim^ cells in the choroid plexus compared to the blood indicates enrichment of CD56^bright^ NK cells in the choroid plexus. Accordingly, we could not detect GrB^+^ NK cells, presumptive CD56^dim^ NK cells, within the choroid plexus stroma from controls, and rarely in MS cases, using multiplex immunohistochemistry. The upregulation of CD54 in CD56^dim^ NK cells in MS was unique to the choroid plexus and differed from CD56^dim^ NK cells in the circulation and the septum. However, the proportion of CD56^dim^ NK cells did not differ between groups. This suggests an altered migratory capacity of CD56^dim^ NK cells within the MS choroid plexus. Although the higher frequency of CD56^bright^ NK cells measured by CyTOF in the choroid plexus from MS donors relative to controls was not significant, multiplex immunohistochemistry confirmed the accumulation of GrK^+^ NK cells in the choroid plexus stroma of MS donors from an independent cohort. Future studies should uncover if CD56^bright^ NK cells continue their journey across the blood-CSF barrier and the ependymal layer lining the periventricular brain. Supporting this idea, the CSF ratio of CD56^bright^/CD56^dim^ NK cells is higher in MS patients than in controls [46, 48], and it is further increased upon treatment with the anti-CD25 antibody therapy daclizumab [5]. In conclusion, circulating CD56^bright^ NK cells in MS may infiltrate into periventricular brain areas directly across the blood-brain barrier, and indirectly via the choroid plexus-CSF route.

The immune landscape of the choroid plexus revealed by this study is compatible with previous descriptions [45, 60]. Comparison of post-mortem blood and choroid plexus samples of the same donors confirmed a strong contribution of blood-derived immune cells in the choroid plexus, which precluded a clear distinction between circulating and stromal immune cells. Because of this technical limitation, we compared our CyTOF data with previous histopathological studies. Myeloid cells are the most abundant immune cells in the choroid plexus, while the high proportion of granulocytes likely originates from the circulation [45, 60]. Likewise, the higher proportion of CD8^+^ relative to CD4^+^ T cells in the choroid plexus could also reflect their aberrant abundance in post-mortem blood [12]; this contribution from the blood to the single-cell suspension may have prevented the detection of choroid-plexus specific changes in the stroma such as an accumulation of CD8^+^ T cells in the choroid plexus from MS donors, previously described by our group using immunohistochemistry [45]. The scarcity of B cells, despite the high vascularisation, is in line with previous reports [45, 60]; while the detection of rare ASCs in the choroid plexus but not in the blood reassures their tissue specificity [60].

Notably, our study with post-mortem blood uncovered a higher frequency of B cells in MS donors relative to controls. While the efficacy of peripheral B cell-depleting therapies suggests a pathogenic role of circulating B cells in MS [20, 36], we could not find evidence in the literature showing an accumulation of B cells in the blood of MS patients, and a recent study showed similar frequencies relative to controls [48]. Although most B cells lack T-bet expression, a small subset of T-bet^+^ B cells shows a trend towards more abundance in MS post-mortem blood, in line with recent literature [10, 59]. Moreover, the expression of T-bet within T-bet^+^ B cells was higher in MS donors relative to controls. T-bet expression on B cells has been associated with increased pathogenic responses [3, 42]. The relevance of T-bet^+^ B cells in MS should be further confirmed in fresh blood samples.

Many MS studies either make use of a healthy control group or study the MS disease course over time within the same patient group. Here, we aimed to uncover MS-specific immune changes by comparing our findings in MS donors to an extra neurological control group consisting of demented cases next to our non-neurological controls. All but one of our demented controls were diagnosed with Alzheimer’s disease (AD). By including this extra control group, we found that the increases in CD56^bright^ NK cells and ASCs were specific to MS compared to both dementia and non-neurological controls. Although our study was not powered for focusing on dementia, we found that the abundance of CD8^+^ T cells in the septal periventricular area was higher compared to the non-neurological controls, following the trend seen in MS donors. The most notable increase was seen in CD8^+^ tissue-resident memory T cells, a population that has been found in the ageing brain before [53]. In AD, higher numbers of circulating CD8^+^ T_EMRA_ cells relative to healthy donors were negatively associated with cognition [13]. In addition, clonally expanded CD8^+^ T cells were more abundant in the CSF of AD patients compared to healthy controls [13]. Although our findings were not statistically significant, they highlight a role for periventricular CD8^+^ T cells in AD pathology. Moreover, we identified an activated phenotype of periventricular B cells in demented controls, consisting of higher expression of CD25 and CD49d. Interestingly, an accumulation of activated B cells was found in the circulation of AD mouse models before, together with infiltration of B cells into the brain parenchyma [25]. Future studies with increased sample size should confirm these results and reveal how T and B cells play a role in AD pathogenesis.

## Conclusions

In sum, our study provides a characterization of the periventricular immune landscape in MS and reveals the involvement of CD56^bright^ NK cells in local MS brain pathology. Moreover, we explored the relative contributions of the choroid plexus and peripheral blood to the immune composition of the periventricular MS brain. We speculate that the migratory phenotype of circulating CD56^bright^ NK cells in MS facilitates entry into the CNS directly through the blood-brain barrier and indirectly across the choroid plexus-CSF route. Our findings highlight the importance of CD56^bright^ NK cells in the CNS and their potential as a therapeutic target in MS. Instead of indiscriminately preventing immune cell surveillance in the CNS, an expansion of immunoregulatory CD56^bright^ NK cells could selectively suppress neuroinflammation while minimizing side effects for patients suffering MS.

## Materials and methods

### Human samples and study design

Fresh post-mortem tissue was obtained from donors by rapid autopsy from the Netherlands Brain Bank. Donors included cases with clinically diagnosed and neuro-pathologically confirmed progressive MS (n = 12), cases with dementia (neurological controls or NC; n = 8) and control cases without neurological or autoimmune diseases (non-neurological controls or NNC; n = 10). Septum pellucidum and choroid plexus from the right lateral ventricle were stored in Hibernate-A medium (Thermo Fisher, #A1247501); and post-mortem blood of all included cases was collected in EDTA-coated tubes (BD Biosciences, #367525). Post-mortem delay was less than 24h and tissues were stored for less than 24h (sample delay) before starting the isolation of immune cells.

Formalin-fixed paraffin-embedded tissue from choroid plexus and periventricular areas was obtained from patients with clinically diagnosed MS (choroid plexus, n=10; periventricular areas, n=7) and non-neurological controls (choroid plexus, n=8; periventricular areas, n=5) by rapid autopsy from the Netherlands Brain Bank and Multiple Sclerosis Society Tissue Bank, funded by the Multiple Sclerosis Society of Great Britain and Northern Ireland, registered charity 207,495). All the MS periventricular area blocks contained lesions.

All parties received permission to perform autopsies, for the use of tissue and access to medical records for research purposes. Relevant clinical information of the donors is summarised in Table 1 and Supplementary Figs. 1a, 4a, 6a.

### Sample processing

The choroid plexus was washed twice with ice-cold PBS and incubated with 1 mM EDTA pH 7.2 in HBSS (Thermo Fisher, #14175095) rotating for 1h at 37°C to loosen the tight junctions of the epithelium, washed again and thoroughly cut with sharp scissors and digested with proteolytic enzymes (2 U/mL Liberase TL, Sigma Aldrich, #5401020001) and DNase (33 µg/mL; Sigma Aldrich, #11284932001) for 30 min at 37°C, resuspending every 10 min. Digestion was stopped with RP-HE (RPMI 1640, Thermo Fisher; 10% FCS (Corning, #35-079-CV), 10 mM EDTA (ThermoFisher, #15575020) 20 mM HEPES (Gibco, #156300-056), 50 µm 2-mercaptoethanol (Gibco, #31350010) for 5 min on ice. The cell suspension was filtered through a 70 µm cell strainer (Life Sciences) to remove undigested pieces of tissue. Immune cells were isolated from this cell suspension using a 70-30% Percoll (Sigma Aldrich, #17-0891-01) gradient as follows. The eluent containing the single-cell suspension was centrifuged, the pellet was resuspended in 70% Percoll, and 30% Percoll was carefully layered on top. Following centrifugation at 900 g for 30 min at 22°C, immune cells in the 70-30% interphase were collected and washed twice with RP10 (RPMI 1640, Thermo Fisher; 10% FCS, 1% penicillin/streptomycin (Invitrogen, #15140122) 1% glutamine (Thermo Fisher, #25030-024) and counted. For the septum, an identical isolation protocol was used leaving out the first step of incubation with EDTA.

Post-mortem blood was diluted with 1% citrate in PBS and filtered through a 70 µm cell strainer to remove clots. Diluted and filtered blood was then carefully layered on top of Lymphoprep (STEMCELL Technologies, #07851). The gradient was centrifuged at 800 g for 30 min at room temperature. The immune cells in the interphase were collected and washed with 1% citrate in PBS. Erythrocytes were lysed with ACK Lysing Buffer (Thermo Fisher, #A1049201) for 5 min at room temperature. Lysis was stopped with 1% citrate in PBS and cells were washed twice with RP10 and counted.

### Viability staining, fixation and freezing of immune cells

Isolated immune cells were washed with HBSS^-/-^ (without Mg^+2^, Ca^+2^ and phenol red, #14175-095). Cells were distributed in wells of a 96-well V-bottom plate (0.5 to 1 million cells/well for septum and choroid plexus cells, 5 million cells/well for blood cells) and washed again with HBSS^-/-^. Cells were washed with Maxpar PBS (Fluidigm, #201058) and stained with the viability marker Cell-ID™ Cisplatin-198Pt (Fluidigm, #201198) for 5 min at 37°C according to the manufacturer’s instructions. Cells were then washed three times with RP10 and fixed with Maxpar Fix I Buffer (Fluidigm, #201065) for 10 min at room temperature. Fixed cells were centrifuged at 800 g for 7 min at 4°C. Cell pellets were resuspended in 10% DMSO in FCS and put in a Mr Frosty Freezing Container at −80°C for 24 h. Samples were stored at −80°C until staining and sample acquisition.

### Antibody labelling and titration

Antibody labelling with the indicated metal tag was performed using the MaxPAR® antibody conjugation kit (Fluidigm) according to the manufacturer’s instructions. Purification of the bound antibody was performed with high-performance liquid chromatography (Thermo Fisher) and subsequently concentrated by filtering with a 10kDa filter (Merck Millipore) in a swing-out bucket at 4000RPM for 15 minutes. The end volume was determined and an equal volume of antibody stabilizer buffer (Fluidigm; supplemented with 0.05% sodium azide) was added before the antibodies were stored at 4°C. All antibodies used in this study were titrated using both fixed and unfixed thawed PBMCs and the most optimal concentrations with the least spillover were chosen. Concentrated antibody cocktails for surface and nuclear antigen detection were made, aliquoted and stored at −80°C, as previously described (Schulz et al, 2019).

### Staining protocol

During the staining, reagents were cooled on ice, centrifugation steps were performed at 800g for 7 min at 4°C with acceleration 9 and deceleration 7, and incubations were performed at room temperature. Samples were thawed rapidly and washed twice with 8 mL of Maxpar® PBS (Fluidigm, #201058). Cell pellets were resuspended in Maxpar® PBS and transferred to a 96 well V-bottom plate (Sigma Aldrich, #M9686). After centrifugation, cells were washed twice with 150 μL of 1X Barcode Perm Buffer (Fluidigm, #201057). Then, samples were incubated with the appropriate barcodes (Fluidigm, #201060) in 1X Barcode Perm Buffer for 30 min, gently mixing after 15 min. After centrifugation, samples were washed twice with 150 μL of cell staining buffer (CSB) (Fluidigm, #201068) and the cells from all samples were pooled. Cells in this combined sample were counted and after centrifugation, cells were incubated with FC block (BioLegend, #422302) diluted in CSB (1:50) for 10 min. Then, the surface antibody cocktail (thawed and centrifuged for 15 minutes at 15,000 × g at 4°C) was diluted with CSB in the appropriate concentration and volume relative to the cell count (Table 2) and added to the combined sample. This was followed by a 30 min incubation, gently mixing after 15 min. After the incubation, cells were washed twice with CSB followed by a wash with Maxpar® PBS. Then, cells were fixed with 1 mL of freshly made 1.6% PFA (Thermo Fisher, #28906) in Maxpar® PBS for 10 min. After centrifugation, cells were permeabilised with a 30 min incubation in 1 mL of FoxP3 Fix/Perm working solution (eBioscience, #00-5523), gently mixing after 15 min. After centrifugation, cells were washed twice with 1X Permeabilisation Buffer (eBioscience, #00-5523). The nuclear antibody cocktail was thawed and centrifuged for 15 min at 15,000 × g at 4°C and cells were incubated with the nuclear antibody cocktail in 1X Permeabilisation Buffer with the appropriate concentration to the cell count. The cells were incubated with the nuclear antibody mix for 45 min, gently mixing every 15 min. Cells were then washed three times with 1X Permeabilisation Buffer and fixed with 1 mL of freshly made 1.6% PFA in Maxpar® PBS for 10 min. After centrifugation, nucleated cells were stained with Maxpar Intercalator (Fluidigm, #201192B) diluted 1:4000 in Maxpar Fix and Perm Buffer (Fluidigm, #201067) overnight at 4°C, until sample acquisition.

### Sample acquisition

Cells in Intercalator solution were washed twice with CSB and divided over approximately 1 × 10^6^ cells per tube, followed by washing with cell acquisition solution (CAS) (Fluidigm, #201241) right before acquisition. Samples were filtered and calibration beads (Fluidigm, #201078) were added to the suspension to 15% of the final volume. Cells were acquired on the Helios™ (Fluidigm), with an event rate of 250-350 events per second. Runs took approximately 30 – 45 minutes. During the day, tuning of the machine was performed during start-up and after 4 hours of sample acquisition. Within each barcoded set of samples, one reference sample was included to monitor possible differences in staining intensity between barcodes due to technical variation in the staining protocol or daily changes in instrument functioning.

### Generation of the reference sample

The reference sample contained peripheral blood mononuclear cells (PBMCs) obtained from the blood of 3 healthy controls of which a part was stimulated with a cytokine cocktail to induce expression of each protein and transcription factor included in the CyTOF panel. Unstimulated and stimulated PBMCs were combined, stained for viability as described earlier, fixated and stored in aliquots at −80°C until further use.

### CyTOF data pre-processing and tissue comparisons

Acquired samples were randomised using Gaussian negative half zero randomization in CyTOF Software version 6.7. The FCS files were normalised using bead normalisation, concatenated and debarcoded using the CyTOF Software version 6.7. The concatenated FCS files were uploaded into FlowJo V10 to perform the clean-up where normalization beads, cell debris and cell doublets were removed using DNA, beads and Gaussian parameters. Next, live cells showing negative reactivity for viability marker Cell-ID™ Cisplatin-198Pt and dim-to-positive reactivity for CD45-89Y were selected and used for the further processing steps. To anticipate similarities between choroid plexus, septum and blood-derived immune cells, 10 000 cells per donor tissue were imputed and visualised in one UMAP (Uniform Manifold Approximation and Projection).

### CyTOF data processing and statistical analysis

Data analysis was performed in R version 4.0.3 (2020-10-10). Clinical data was analysed by the Kruskal-Wallis rank-sum test followed by the Wilcoxon rank-sum test with continuity correction. Quantitative data are shown as independent data points in box plots indicating the median and interquartile range. CyTOF data was analysed following a published workflow [38] (update 28 April 2020) with some modifications. Marker expression data was arcsinh transformed with a cofactor of 5. We decided to not normalise the data after visually confirming that the expression pattern distribution did not differ between batches of stain/run days (Supplementary Fig. 8).

Clustering of the single-cell data was performed with FlowSOM [60] and ConsensusClusterPlus [65], using the lineage markers labelled as “type” (Table 2). We clustered each tissue separately and pooled the samples from all donors; we subsampled the data using the 75^th^ percentile of the number of cells per tissue as a threshold, to prevent big samples from driving the clustering (see Supplementary Figs. 1b, 4b, 6b). The metaclusters obtained (20 for the main populations, 12 for the T cell lineage and 8 for the NK and B cell lineages) were manually annotated and merged based on visualisation using heatmaps with normalised median marker expression per cluster and dimensionality reduction plots (tSNE and UMAP).

We used a generalised linear mixed model (GLMM) for analysing the differential abundance of cell populations and a linear mixed model (LMM) for the differential expression of “state” phenotypic markers per cell population. The models were built through the R package *diffcyt*, with a few modifications. The GLMM response variable is the cell counts per cell type (those with more than 100 cells) and sample, and the explanatory variable is the disease group, as a fixed effect. To model the overdispersion in proportion estimates (uncertainty is higher when proportions are calculated from samples with a low total number of cells), we included the sample ID as a random effect. The LMM response variable is the median marker expression of the state markers per cell type (those with more than 100 cells) and sample. Differential expression of markers was only considered in cell populations for which they were biologically meaningful (for example, FoxP3 was only considered in T and B cells), and in which the expression values were higher than 0 in at least a third of the samples. Weights were assigned to each cluster and sample based on the number of cells, to account for differences in uncertainty in the calculation of the medians. We specified the contrasts MS vs. NNC, MS vs. NC and NNC vs NC. We corrected the p-values for multiple testing with the Benjamini-Hochberg procedure, with a false discovery rate (FDR) cut-off of 10%; for differential abundance, we corrected for cell population per tissue type, and for differential expression, we corrected for state marker per cell population.

Correlations were performed using the functions *rcorr* and *corr.test*, as implemented in the packages *psych* and *Hmisc*, respectively, using the Benjamini-Hochberg method to adjust p-values. Only patients with samples analysed in all three tissues were used. Significant correlations (adjusted P ≤ 0.1) were visualised in a network plot using Cytoscape. The strength of the correlation was calculated as −log10(adjusted P) and shown as the width of edges.

### Multiplex immunohistochemistry

Sections of 5 µm-thickness from formalin-fixed paraffin-embedded tissue were cut with a microtome and mounted on SuperFrost Plus slides. Sections were deparaffinised in xylene and rehydrated with a series of graded ethanol (100%, 90%, 80% and 70% for 2 min). Antigen retrieval was performed in citrate buffer pH 6.0 at 95°C for 30 min in a water bath. Endogenous peroxidases were blocked using 0.3% H_2_O_2_ in PBS for 15 min. Slides were blocked using 1% bovine serum albumin (BSA) in PBS with 0.05% Tween-20 for 30 min. Sections were then incubated with a single unconjugated primary antibody (Table 3) in 10x diluted blocking buffer for either 60min at room temperature or 24h at 4°C. Slides were washed in PBS-Tween-20 and incubated with Envision+ Dual link HRP (Dako, #K4061) for 30 min. Sections were then incubated with the respective Opal fluorochrome (Opal 480, Opal 520, Opal 570, Opal 620, Opal 650, Opal 780) at a 1:250 dilution made in tyramide signal amplification reagent (Akoya, #FP1498) for 60 min. Slides were washed in PBS-Tween-20 and a heating step of citrate buffer pH 6.0 at 98°C for 30 min in a water bath was performed for primary antibody removal. Afterwards, the tissue was stained by repeating staining cycles in series as described above with a primary antibody removal step in each cycle. Finally, slides were counterstained with DAPI (Invitrogen, #D1306) for 5 min and mounted with Prolong Gold (Invitrogen, #P36930). Slides were imaged using the Vectra 3.0 spectral imaging system (PerkinElmer), with a low magnification scanning at 10x to get an overview of the slide. To get representative images of the tissue section, a minimum of 2 regions of interest across the fields were chosen and scanned at 40x magnification. Spectral unmixing was performed using InForm advanced image analysis software (PerkinElmer) and image segmentation was done in Nis Elements (Nikon). In short, nuclei detection was based on DAPI signal, and a mask was made from nuclei that were directly surrounded by CD45 signal. This mask was used to further select NK cells based on NKp46 signal. The NK cell mask was then divided based on granzyme K and granzyme B expression. Cells within the lumen of vessels from the choroid plexus sections were excluded manually. All cell counts were then normalised to the total tissue area. If data was normally distributed as assessed by the Shapiro-Wilk test, an ANOVA test or unpaired t-test was used; if the data did not pass normality, Kruskal-Wallis test or Mann-Whitney test was used. Staining, imaging and analysis were performed blinded.

## Acknowledgements

We would like to acknowledge all the donors and their families who made this study possible, as well as the Netherlands Brain Bank, and especially Michiel Kooreman for his support in the collection of the human samples. We thank the Microscopy and Cytometry Core Facility from the Amsterdam UMC for excellent technical support, especially Juan J. Garcia Vallejo and Cora Chadick. We appreciate the inspiring discussions with the members of the Molecular Cell Biology and Immunology department, especially Mike de Kok, Jan Verhoeff and Reina Mebius. We are grateful to Marvin M. van Luijn for his valuable input on T-bet^+^ B cells. Stefanos Prouskas kindly provided information on the lesion location for immunohistochemical validation. We appreciate the support on panel design from Olga Karpus, on analysis from Sofie van Gassen, and on statistical analysis from Mark van de Wiel. This work was funded by the pilot grant 20-1087 MS from the Dutch MS Research Foundation to SRL, GK and HEV.

## Author contributions

LvO and SRL conceived the study and designed the CyTOF experiments, optimised the immune cell isolation. SRL, LvO, CRM, SvP and AK performed the isolations and stainings for CyTOF. LvO and SvP performed the CyTOF measurements. SRL analysed the CyTOF data and prepared the figures with input from LvO, CRM and ER. AK, with the help of CRM, designed, performed and analysed the multiplex immunohistochemical validation. SRL, LvO, CRM and AK interpreted the data and wrote the manuscript. GK and HEV provided funding and HEV supervised the study. All authors revised the manuscript.

## Competing interests

The authors declare that they have no competing interests

## Supplementary figures

**Supplementary Fig. 1.**
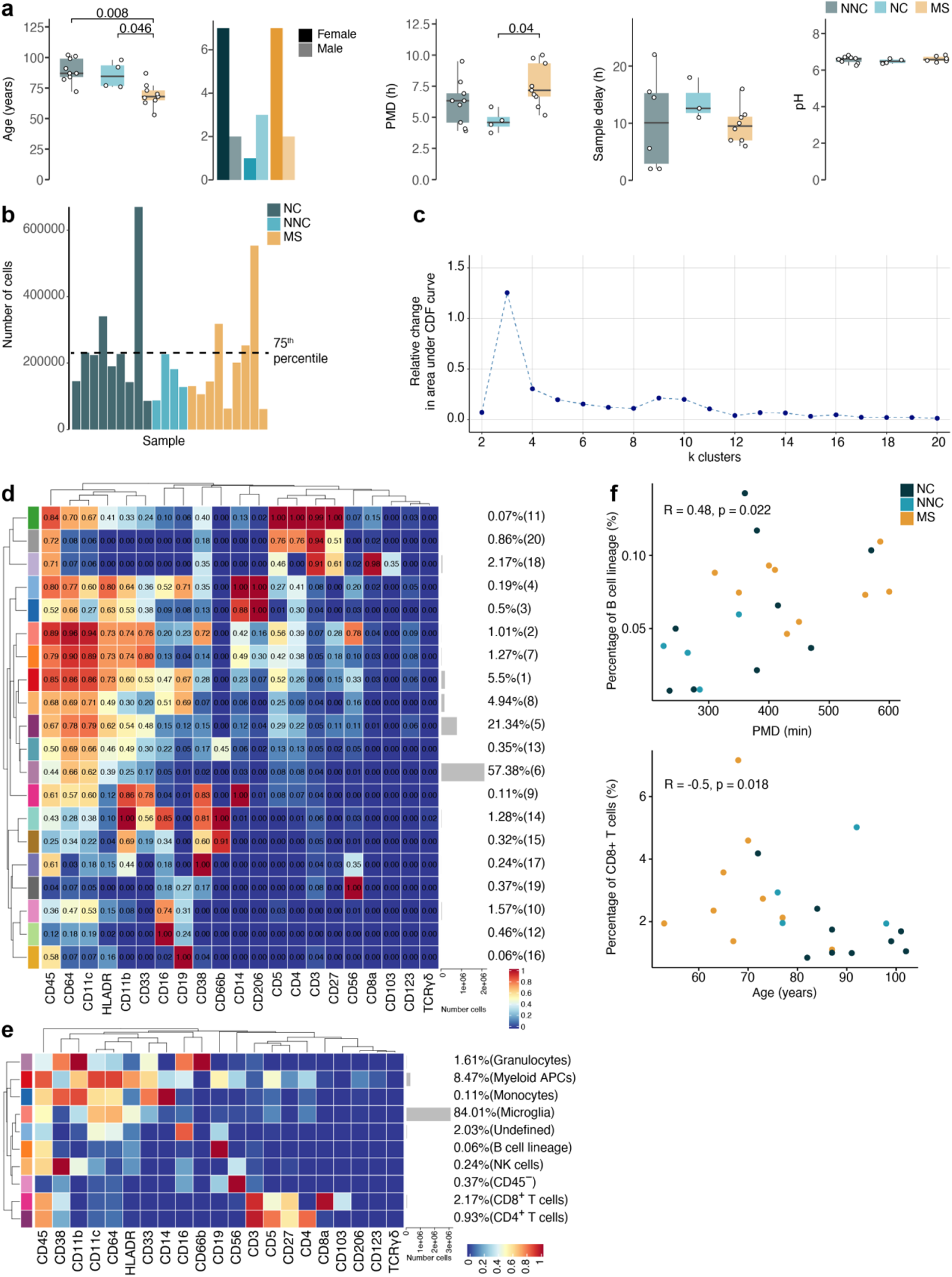
Immune phenotyping of the septum using mass cytometry reveals inflammation in multiple sclerosis involving T cells and natural killer cells. **(a)** Clinical parameters of the septum samples. Age and PMD (post-mortem delay): pairwise comparisons using Wilcoxon rank-sum test with continuity correction; p-values adjusted with Benjamini-Hochberg method. Sample delay and pH: Kruskal-Wallis rank-sum test. **(b)** Bar plot showing the number of cells used for clustering from each septum sample. A threshold was set at the 75th percentile to avoid over-representation of certain samples in the analysis. **(c)** Delta area (“elbow”) plot indicating the relative increase in cluster stability when clustering into k groups the 100 SOM clusters generated by FlowSOM. CDF: Consensus Cumulative Distribution Function. **(d)** Median marker intensities of the “type” markers across septum-derived cell populations obtained with FlowSOM and ConsensusClusterPlus. The colour in the heatmap represents the median of the arcsinh, 0-1 transformed marker expression calculated over cells from all samples. The left dendrogram is based on hierarchical clustering with average linkage and Euclidean distance between the 20 metaclusters. **(e)** Median marker intensities of the “type” markers across septum-derived cell populations. **(f)** Dot plots show the significant correlations (P < 0.05) between the percentage of the septum-derived immune cells and sample parameters. Non-normally distributed data: Spearman’s rank correlation rho; normally distributed data: Pearson’s correlation. **(d, e)** Horizontal grey bars show the percentage out of the total cells.

**Supplementary Fig. 2.**
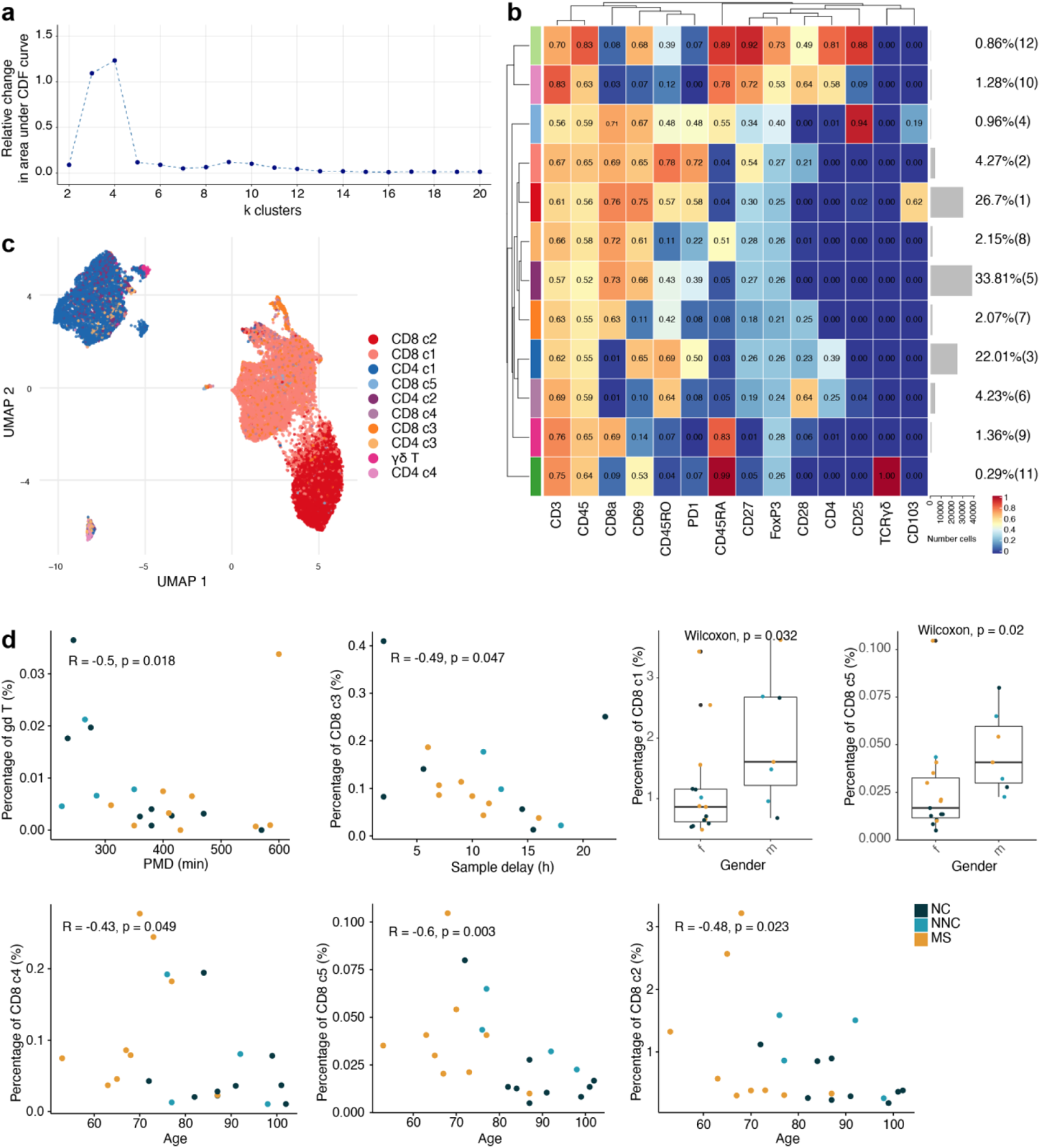
Phenotyping the T cell populations in the septum using mass cytometry. Delta area (“elbow”) plot indicating the relative increase in cluster stability when clustering into k groups the 100 SOM clusters generated by FlowSOM. CDF: Consensus Cumulative Distribution Function. **(b)** Median marker intensities the “type” markers across T cell lineage populations of the septum obtained with FlowSOM and ConsensusClusterPlus. Horizontal grey bars show the percentage out of the total cells. The colour in the heatmap represents the median of the arcsinh, 0-1 transformed marker expression calculated over cells from all the samples. The left dendrogram is based on the hierarchical clustering with average linkage and Euclidean distance between the clusters. **(c)** UMAP plot based on the arcsinh-transformed expression of the 14 “type” markers in the T cell subsets in the septum. A subset of 1,000 randomly selected cells per sample is shown, coloured according to the manually annotated clusters. **(d)** Dot plots show the significant correlations (P < 0.05) found between the percentage of the T cell subsets of the septum and clinical sample parameters. Spearman’s rank correlation rho is shown for non-normally distributed data and Pearson’s correlation for normally distributed data. Boxplots show the percentage of cells in females (f) and males (m).

**Supplementary Fig. 3.**
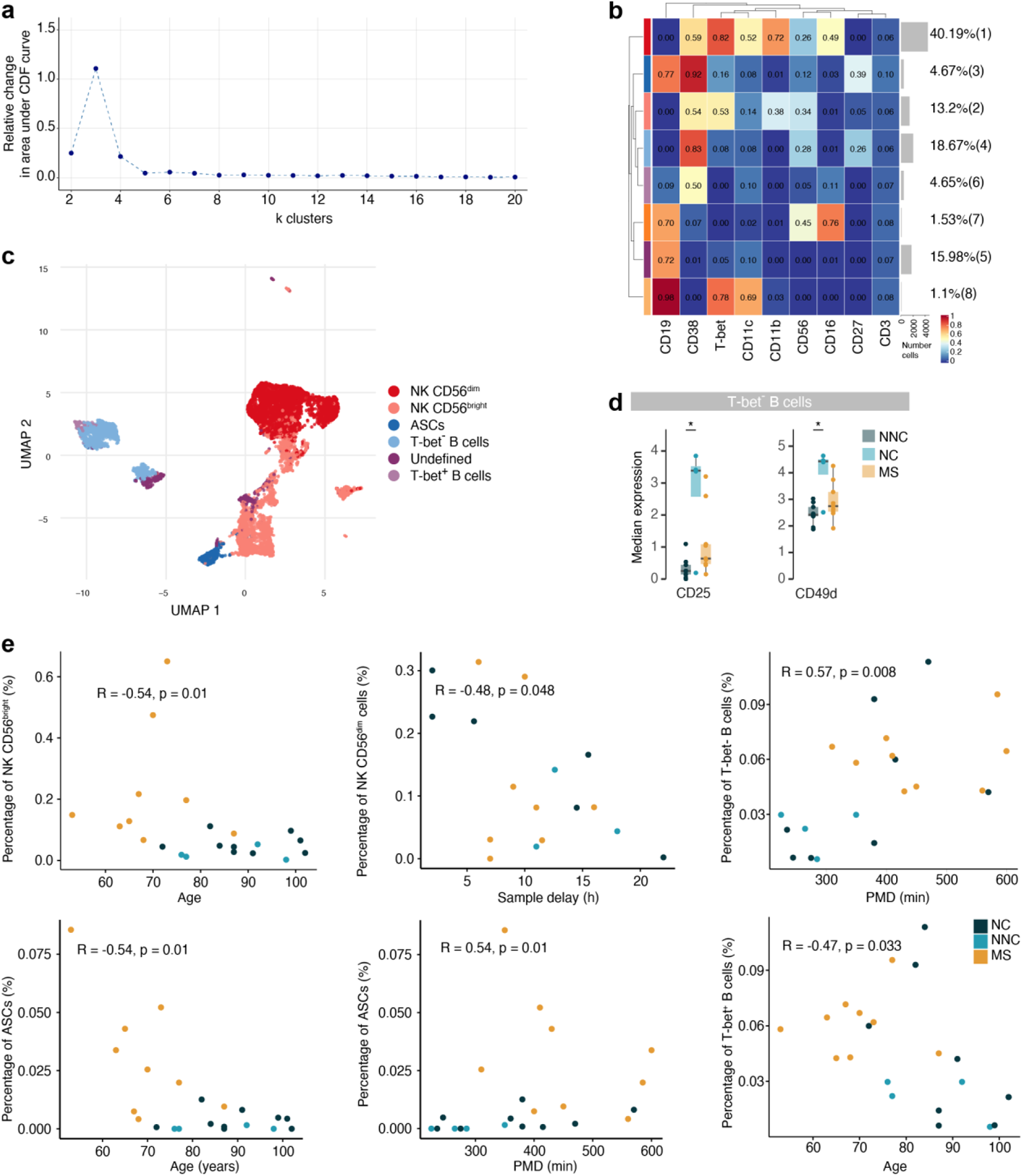
NK & B cell subpopulations in the septum. **(a)** Delta area (“elbow”) plot indicating the relative increase in cluster stability when clustering into k groups the 100 SOM clusters generated by FlowSOM in the NK & B cell populations. CDF: Consensus Cumulative Distribution Function. **(b)** Median scaled intensities of the “type” markers across NK & B cell lineage populations of the septum obtained with FlowSOM and ConsensusClusterPlus. Horizontal grey bars show the percentage out of the total cells. The colour in the heatmap represents the median of the arcsinh, 0-1 transformed marker expression calculated over cells from all the samples. The left dendrogram is based on the hierarchical clustering with average linkage and Euclidean distance between metaclusters. **(c)** UMAP plot based on the arcsinh-transformed expression the “type” markers in the NK & B cell lineage populations of the septum. A subset of 1,000 randomly selected cells is shown, coloured according to the manually annotated clusters. **(d)** Boxplots display the median marker expression of CD25 and CD49d in T-bet^-^ B cells from the septum. *adjusted P < 0.1. **(e)** Dot plots show the significant correlations (P < 0.05) found between the percentage of the septum NK & B cell subsets and clinical sample parameters. Spearman’s rank correlation rho is shown for non-normally distributed data and Pearson’s correlation for normally distributed data. NNC: Non-neurological controls; NC: neurological controls; MS: multiple sclerosis.

**Supplementary Fig. 4.**
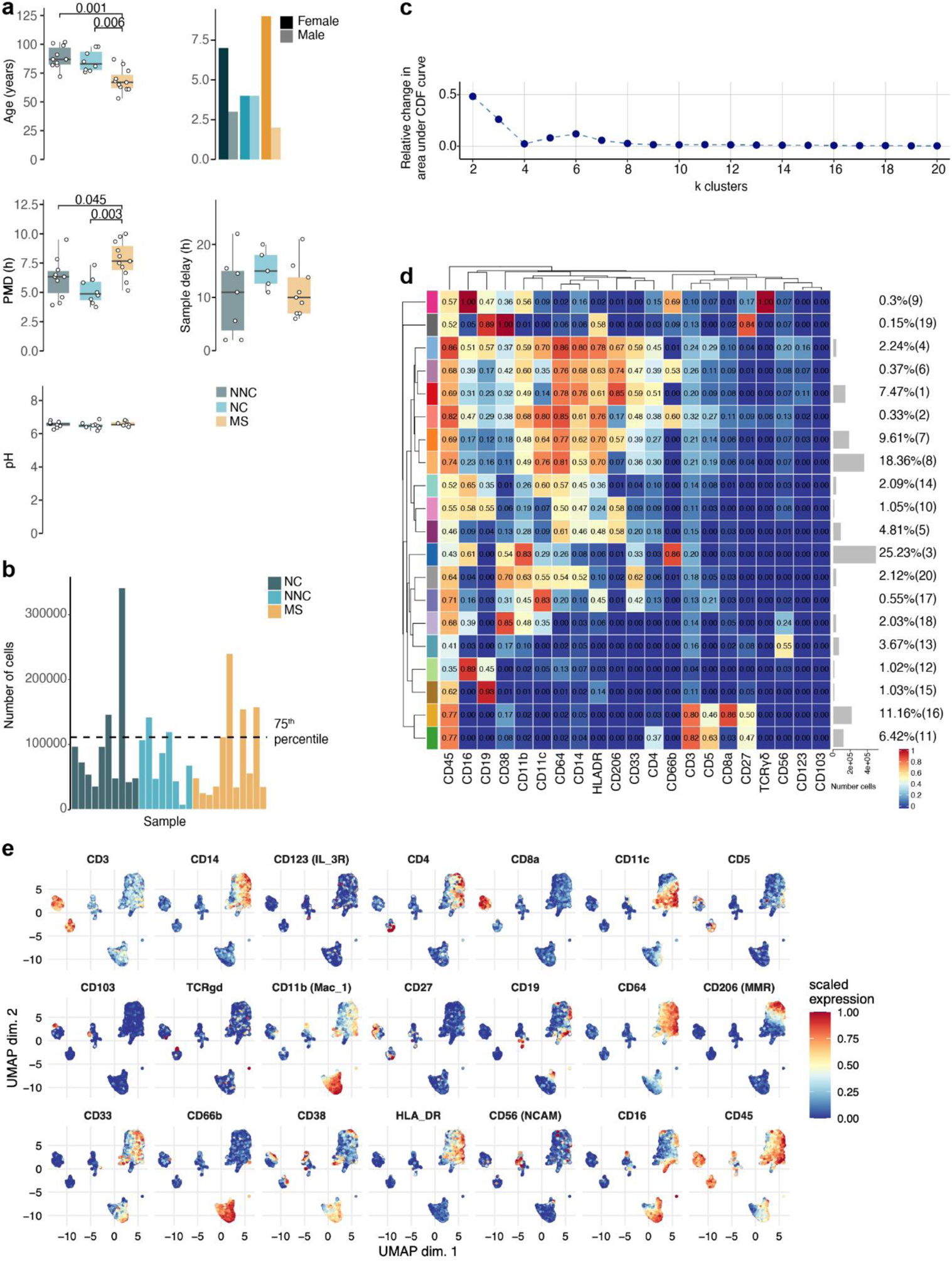
Immune phenotyping of the choroid plexus using mass cytometry. **(a)** Clinical parameters of the choroid plexus samples. Age and PMD (post-mortem delay): pairwise comparisons using Wilcoxon rank-sum test with continuity correction; p-values adjusted with Benjamini-Hochberg method. Sample delay and pH: Kruskal-Wallis rank-sum test. **(b)** Bar plot of the number of cells used for clustering from each choroid plexus sample. A threshold was set at the 75th percentile to avoid over-representation of certain samples. **(c)** Delta area (“elbow”) plot indicating the relative increase in cluster stability when clustering into k groups the 100 SOM clusters generated by FlowSOM. CDF: Consensus Cumulative Distribution Function. **(d)** Median scaled intensities of the “type” markers across choroid plexus-derived cell populations obtained with FlowSOM and ConsensusClusterPlus. Horizontal grey bars show the percentage out of the total cells. The colour in the heatmap represents the median of the arcsinh, 0-1 transformed marker expression calculated over cells from all the samples. The left dendrogram is based on the hierarchical clustering with average linkage and Euclidean distance between the 20 metaclusters. **(e)** UMAP plots based on the arcsinh-transformed expression of the “type” markers in the choroid plexus-derived immune cells. A subset of 1,000 randomly selected cells per sample is shown, coloured according to the expression level of each marker.

**Supplementary Fig. 5.**
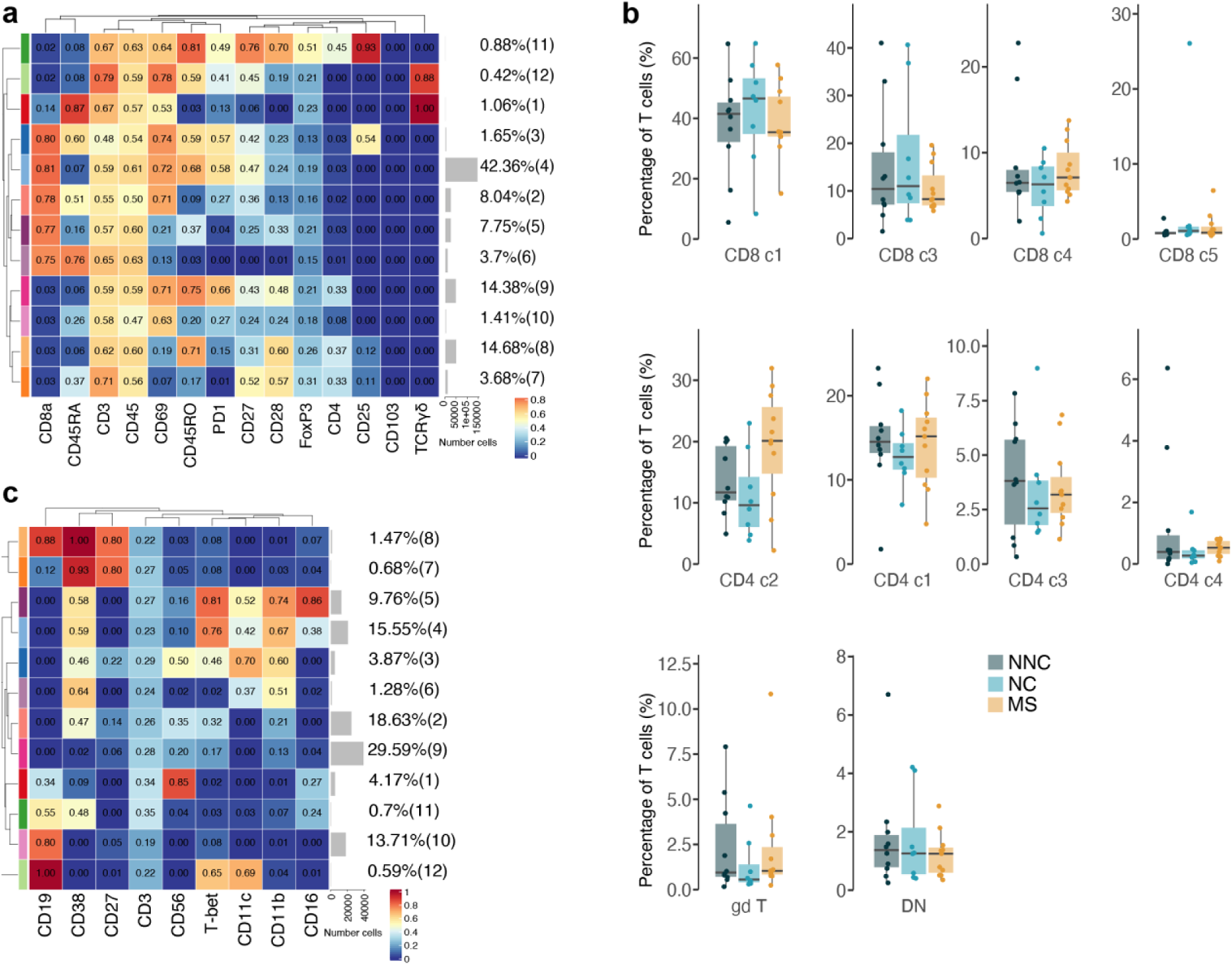
Immune phenotyping of the T cell, NK and B cell subsets in choroid plexus. **(a)** Median scaled intensities of the “type” markers across T cell clusters of the choroid plexus obtained with FlowSOM and ConsensusClusterPlus. **(b)** Percentage of each annotated cell population out of the total number of T cells (CD4^+^ and CD8^+^) from the choroid plexus of NNC, NC and MS donors. **(c)** Median scaled intensities of the “type” markers across NK & B cell lineage populations of the choroid plexus obtained with FlowSOM and ConsensusClusterPlus. **(a, c)** Horizontal grey bars show the percentage out of the total cells. The colour in the heatmap represents the median of the arcsinh, 0-1 transformed marker expression calculated over cells from all the samples. The left dendrogram is based on the hierarchical clustering with average linkage and Euclidean distance between the clusters. NNC: non-neurological controls; NC: neurological controls; MS: multiple sclerosis

**Supplementary Fig. 6.**
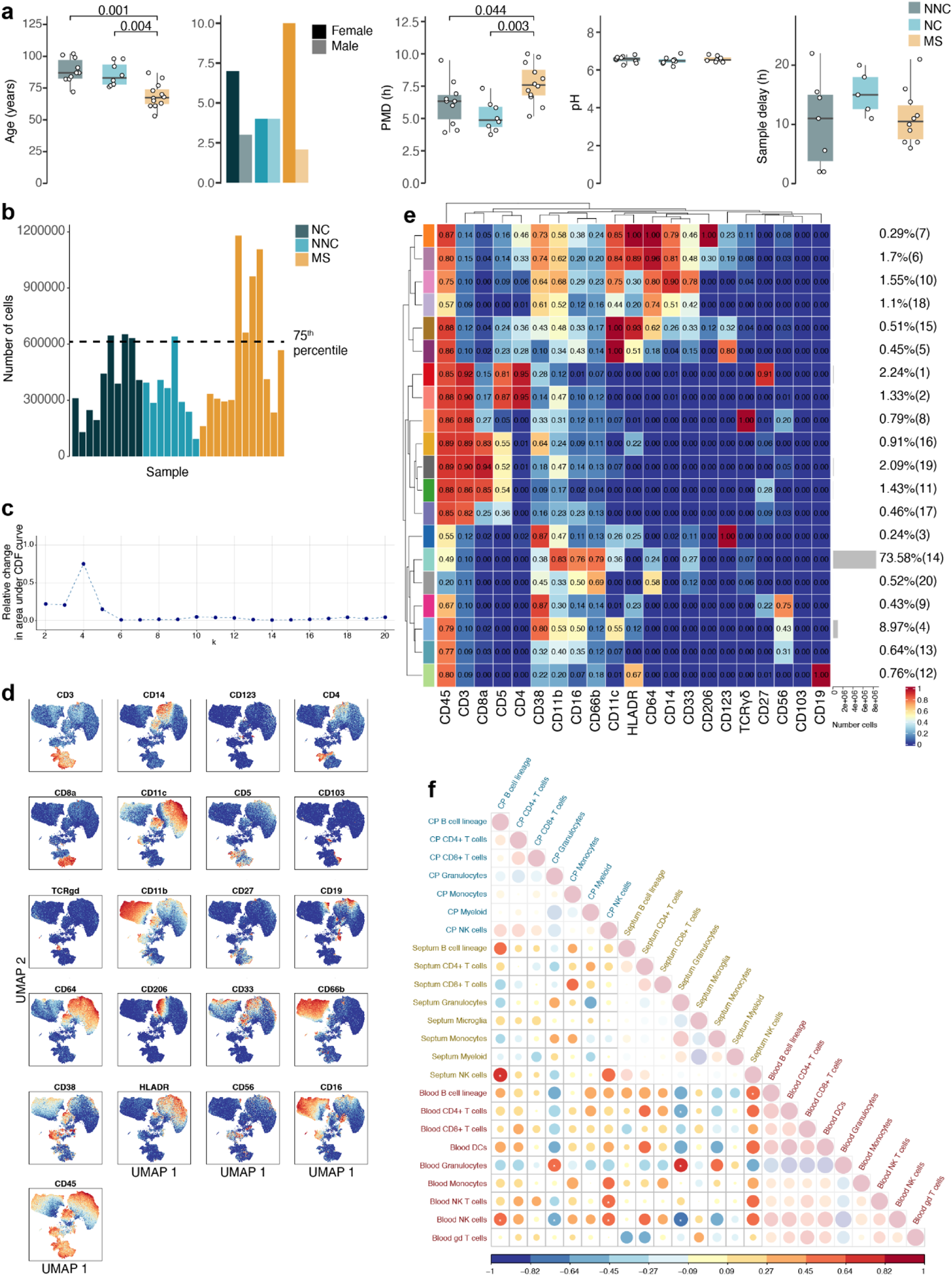
Immune phenotyping of the main immune cell populations in blood and their correlations with those in septum and choroid plexus. **(a)** Clinical parameters of the blood samples. Age and PMD (post-mortem delay): pairwise comparisons using Wilcoxon rank-sum test with continuity correction; p-values adjusted with Benjamini-Hochberg method. Sample delay and pH: Kruskal-Wallis rank-sum test. **(b)** Bar plot of the number of cells used for clustering from each blood sample. A threshold was set at the 75th percentile to avoid over-representation of certain samples. **(c)** Delta area (“elbow”) plot indicating the relative increase in cluster stability when clustering into k groups the 100 SOM clusters generated by FlowSOM. CDF: Consensus Cumulative Distribution Function. **(d)** UMAP plot based on the arcsinh-transformed expression of all markers in the immune cells of the septum, choroid plexus and blood. 10,0000 cells were randomly selected per sample, coloured according to the expression level of each marker. **(e)** Median scaled intensities of the “type” markers across blood-derived cell populations obtained with FlowSOM and ConsensusClusterPlus. Horizontal grey bars show the percentage out of the total cells. The colour in the heatmap represents the median of the arcsinh, 0-1 transformed marker expression calculated over cells from all the samples. The left dendrogram is based on the hierarchical clustering with average linkage and Euclidean distance between the 20 metaclusters. **(f)** Spearman correlation matrix of the proportion of the main immune cell populations among the studied tissues. Positive correlations are displayed in red and negative correlations in blue. The size of the circle is proportional to the correlation coefficients. P-values adjusted with Benjamini-Hochberg method, excluding intra-tissue correlations. * adjusted P < 0.1.

**Supplementary Fig. 7.**
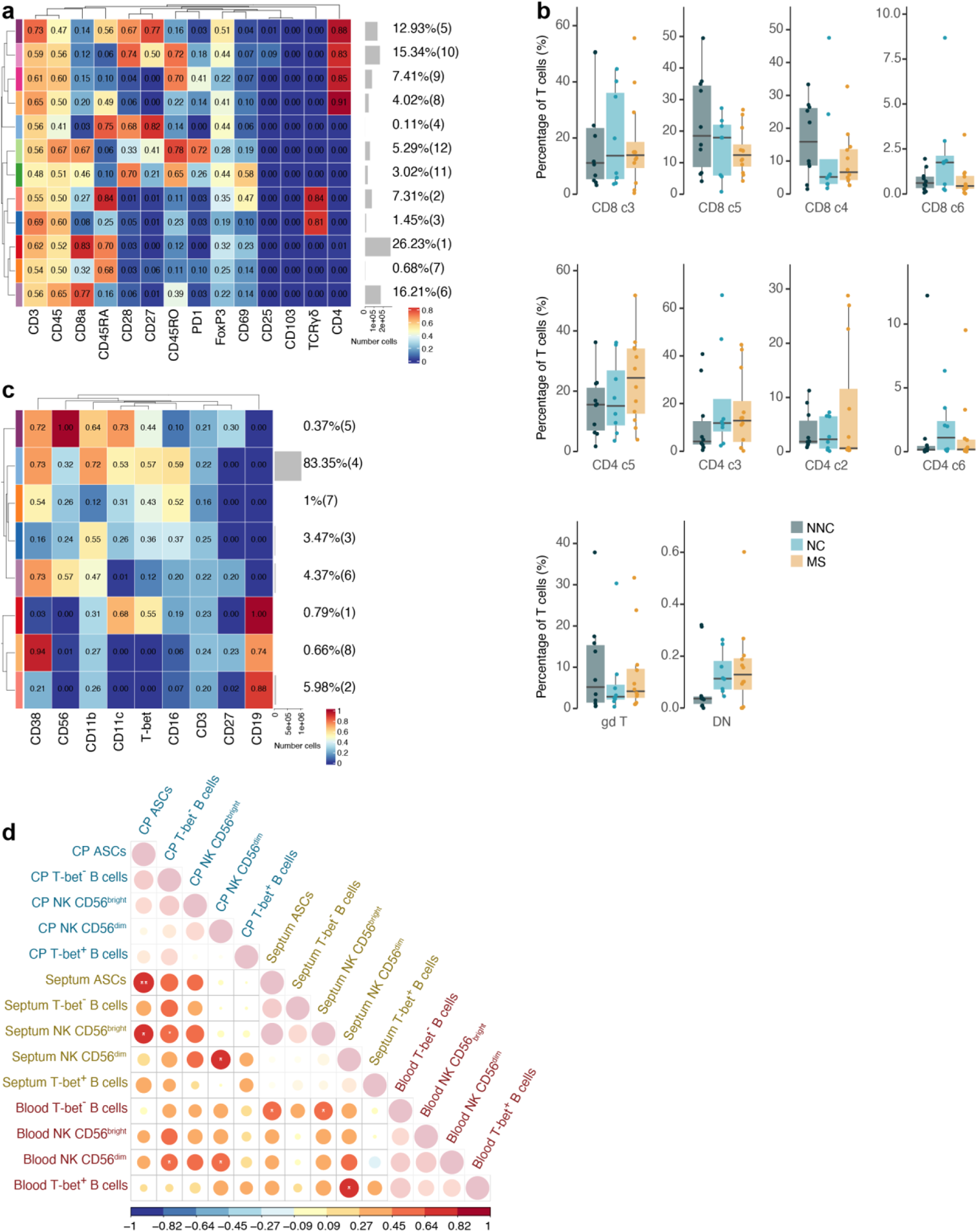
Immune phenotyping of the T cell, NK and B cell subsets in blood and their correlations with those in septum and choroid plexus. **(a)** Median scaled intensities of the “type” markers across T cell clusters in the blood obtained with FlowSOM and ConsensusClusterPlus. **(b)** Percentage of each annotated cell population out of the total number of T cells (CD4^+^, CD8^+^ and γδ T) from the septum of NNC, NC and MS donors. **(c)** Median scaled intensities of the “type” markers across NK & B cell lineage populations in the blood obtained with FlowSOM and ConsensusClusterPlus. **(d)** Spearman correlation matrix of the proportion of NK & B cell populations among the studied tissues. Positive correlations are displayed in red and negative correlations in blue. The size of the circle is proportional to the correlation coefficients. P-values adjusted with Benjamini-Hochberg method, excluding intra-tissue correlations. * adjusted P < 0.1. NNC: non-neurological controls; NC: neurological controls; MS: multiple sclerosis **(a, b)** Horizontal grey bars show the percentage out of the total cells. The colour in the heatmap represents the median of the arcsinh, 0-1 transformed marker expression calculated over cells from all the samples. The left dendrogram is based on the hierarchical clustering with average linkage and Euclidean distance between the clusters.

**Supplementary Fig. 8.**
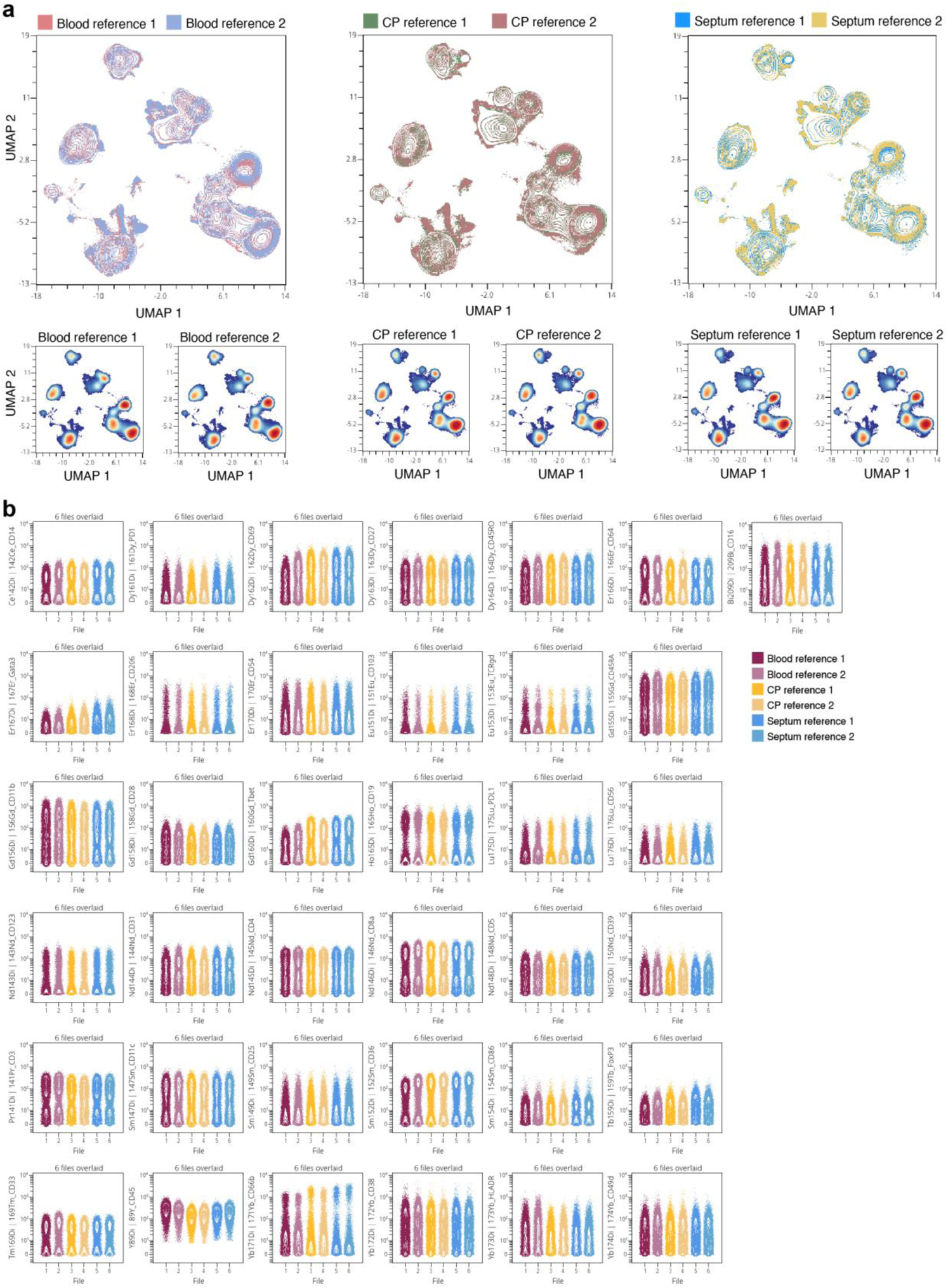
Minor variation in staining intensity between the different barcoded CyTOF runs displayed using a reference sample. **(a)** UMAP plots based on the arcsinh-transformed expression of all markers in the reference sample that were stained together with barcoded runs of septum, choroid plexus and blood samples. 150,000 random cells were selected per sample. The first plot shows an overlay figure where the reference sample is coloured by its barcoded run. The plots below are coloured according to the density. **(b)** Contour plots with 0,1 outlier percentile showing the staining intensity per marker of the reference sample used at different barcodes. The six aliquots of reference samples used were derived from the same sample that was fixated, aliquot and frozen at −80C before. CP: choroid plexus.

**Table S1.** Information of the donors used in this study

| Group | Sample | Gender | Age of death | PMD (h:m) | pH (CSF) | Diagnosis | Disease duration | Braak | Cause of death | CP | Septum | Blood |
| --- | --- | --- | --- | --- | --- | --- | --- | --- | --- | --- | --- | --- |
| NNC | 2018-135 | f | 81 | 4:05 | 6.46 | AD | 5 | 5 | Dementia | x |  | x |
| NNC | 2019-010 | f | 76 | 3:45 | 6.55 | AD | 3 | 5 | Euthanasia | x | x | x |
| NNC | 2019-030 | m | 77 | 4:25 | 6.63 | AD | na | 4 | AD; palliative sedation | x | x | x |
| NNC | 2019-032 | m | 92 | 5:50 | 6.35 | AD | 3 | 4 | Dementia, gastro-intestinal bleeding | x | x | x |
| NNC | 2019-040 | f | 98 | 7:20 | 6.18 | AD | na | 4 | Old age, dehydration due to ceased oral intake | x |  | x |
| NNC | 2020-013 | m | 98 | 4:45 | 6.4 | AD | 10 | na | Pneumonia | x | x | x |
| NNC | 2020-027 | m | 78 | 5:00 | 6.5 | AD | na | na | AD | x |  | x |
| NNC | 2020-036 | f | 85 | 6:05 | 6.88 | dementia | na | na | Carcinoma | x |  | x |
| MS | 2018-107 | f | 83 | 7:40 | 6.54 | PPMS | 34 | na | Ovarian cancer | x |  | x |
| MS | 2019-011 | m | 63 | 10 | 6.52 | PMS | 30 | na | Aspiration pneumonia and sepsis | x | x | x |
| MS | 2019-016 | f | 77 | 9:45 | na | SPMS | >39 | na | Pneumonia | x | x | x |
| MS | 2019-027 | f | 68 | 9:20 | na | SPMS | 18 | na | Infection | x | x | x |
| MS | 2019-031 | f | 53 | 5:50 | 6.76 | PPMS | 17 | na | Euthanasia | x | x | x |
| MS | 2019-043 | f | 61 | 8:05 | 6.41 | SPMS | 22 | na | Sepsis | x |  | x |
| MS | 2019-047 | m | 70 | 5:10 | 6.82 | SPMS | 40 | na | Palliative sedation | x | x | x |
| MS | 2019-069 | f | 67 | 6:40 | 13.12 | SPMS | 39 | na | Sepsis | x | x |  |
| MS | 2019-107 | f | 87 | 7:30 | 6.5 | MS | 31 | na | Ceased oral intake | x | x | x |
| MS | 2019-109 | f | 65 | 7:10 | 6.63 | PPMS | 3 | na | Euthanasia | x | x | x |
| MS | 2019-110 | f | 73 | 6:50 | 6.42 | SPMS | 41 | na | Euthanasia |  | x | x |
| MS | 2020-022 | f | 61 | 8:35 | na | MS | na | na | Euthanasia | x |  | x |
| NNC | 2018-120 | m | 72 | 6:55 | 6.84 | Non-demented control | na | 4 | Myeloma | x | x | x |
| NNC | 2018-123 | f | 82 | 6:20 | na | Non-demented control | na | 3 | Pulmonary disease | x | x | x |
| NNC | 2019-029 | m | 87 | 6:20 | 6.47 | Non-demented control | na | 3 | Liver failure | x | x | x |
| NNC | 2019-042 | m | 82 | 6:30 | 6.53 | Non-demented control | na | 2 | Cancer | x |  | x |
| NNC | 2019-054 | f | 101 | 6:00 | 6.71 | Non-demented control | na | na | Heart failure | x | x | x |
| NNC | 2019-077 | f | 91 | 9:30 | 8.38 | Non-demented control | na | 3 | Old age | x | x | x |
| NNC | 2019-079 | f | 84 | 7:50 | 6.59 | Non-demented control | na | 3 | Cancer | x | x | x |
| NNC | 2019-125 | f | 102 | 3:55 | 6.25 | Non-demented control | na | na | Ileus | x | x | x |
| NNC | 2020-003 | f | 87 | 4:35 | 6.64 | Non-demented control | na | na | Pneumonia in lung fibrosis | x | x | x |
| NNC | 2020-028 | f | 99 | 4:05 | 6.67 | Non-demented control | na | na | Euthanasia | x | x | x |

Multiplex immunohistochemistry
| Group | Sample | Gender | Age of death | PMD (h:m) | pH (CSF) | Diagnosis | Disease duration | Braak | Cause of death | CP | Periventricular areas |
| --- | --- | --- | --- | --- | --- | --- | --- | --- | --- | --- | --- |
| NNC | 2005-034 | m | 56 | 14:00 | 7.03 | Non-demented control | na | 0 | Heart failure | x |  |
| NNC | 2002-018 | f | 92 | 7:00 | 6.45 | Non-demented control | na | 1B | Urosepsis | x |  |
| NNC | 2014-029 | f | 78 | 7:10 | 6.32 | Non-demented control | na | 1A | na | x |  |
| NNC | 2014-043 | f | 60 | 8:10 | 6.58 | Non-demented control | na | 0 | na | x |  |
| NNC | 2014-053 | f | 80 | 7:04 | 6.2 | Non-demented control | na | 0 | na | x |  |
| NNC | 2015-033 | m | 93 | 7:40 | 6.2 | Non-demented control | na | 0A | na | x |  |
| NNC | 2015-035 | m | 73 | 8:00 | 5.37 | Non-demented control | na | 2A | na | x |  |
| NNC | 2018-112 | f | 95 | 4:21 | 6.59 | Non-demented control | na | na | na | x |  |
| NNC | ANW797 | m | 59 | 8:00 | na | Non-demented control | na |  | Euthanasia |  | x |
| NNC | ANW945 | f | 59 | 8:10 | na | Non-demented control | na |  | Euthanasia |  | x |
| NNC | ANW019 | m | 68 | 8:40 | na | Non-demented control | na |  | Euthanasia |  | x |
| NNC | ANW782 | f | 69 | 13:00 | na | Non-demented control | na |  | Pulmonary embolism |  | x |
| NNC | ANW999 | m | 74 | 10:20 | na | Non-demented control | na |  | Euthanasia |  | x |
| MS | 2014-058 | f | 74 | 7:50 | 6.40 | SPMS | 50 | na | Euthanasia | x |  |
| MS | 2014-072 | m | 54 | 7:50 | 6.60 | SPMS | 21 | na | Euthanasia | x |  |
| MS | 2015-006 | f | 57 | 10:40 | 6.76 | SPMS | 25 | na | Euthanasia | x |  |
| MS | 2014-056 | m | 70 | 6:55 | 6.51 | SPMS | 37 | na | Acute heart failure; | x |  |
| MS | 2015-022 | m | 82 | 8:05 | 6.70 | PPMS | 44 | na | Pneumonia | x |  |
| MS | 2018-107 | f | 83 | 7:40 | 6.54 | PPMS | 34 | na | Ovarian cancer | x |  |
| MS | 2018-103 | m | 75 | 9:10 | na | SPMS | na | na | na | x |  |
| MS | 2019-011 | m | 63 | 10 | 6.52 | PMS | 30 | na | Aspiration pneumonia and sepsis | x |  |
| MS | 2019-043 | f | 61 | 8:04 | 6.41 | SPMS | 32 | na | Urosepsis and hydronephrosis | x |  |
| MS | 2019-047 | m | 70 | 5:10 | 6.82 | SPMS | 40 | na | Dehydration, decompensation cordis, MS; palliative sedation | x |  |
| MS | 13-081 (vu16) | m | 66 | 10:55 | 7.28 | PPMS | na | na | Euthanasia |  | x |
| MS | 2019-014 | m | 65 | 7:15 | 6.1 | MS | na | na | Liver cirrhosis |  | x |
| MS | 2018-116 | f | 66 | 9:30 | 6.7 | SPMS | na | na | Euthanasia |  | x |
| MS | 2018-047 | m | 70 | 9:25 | 7.22 | SPMS | na | na | Euthanasia |  | x |
| MS | 2018-046 | f | 51 | 9:10 | 6.87 | SPMS | na | na | Euthanasia |  | x |
| MS | 2018-085 | f | 67 | 11:25 | 6.8 | MS | na | na | Pneumonia |  | x |
| MS | 2018-115 | m | 56 | 6:15 | 6.6 | PPMS | na | na | Respiratory insufficiency |  | x |

PMD, post-mortem delay; h:m, hours:minutes; CP, choroid plexus; f, female; m, male; na, not available; AD, Alzheimer's Disease; MS, multiple sclerosis; SPMS, secondary progressive MS; PMS, progressive MS; PPMS, primary progressive MS.

**Table S2.**
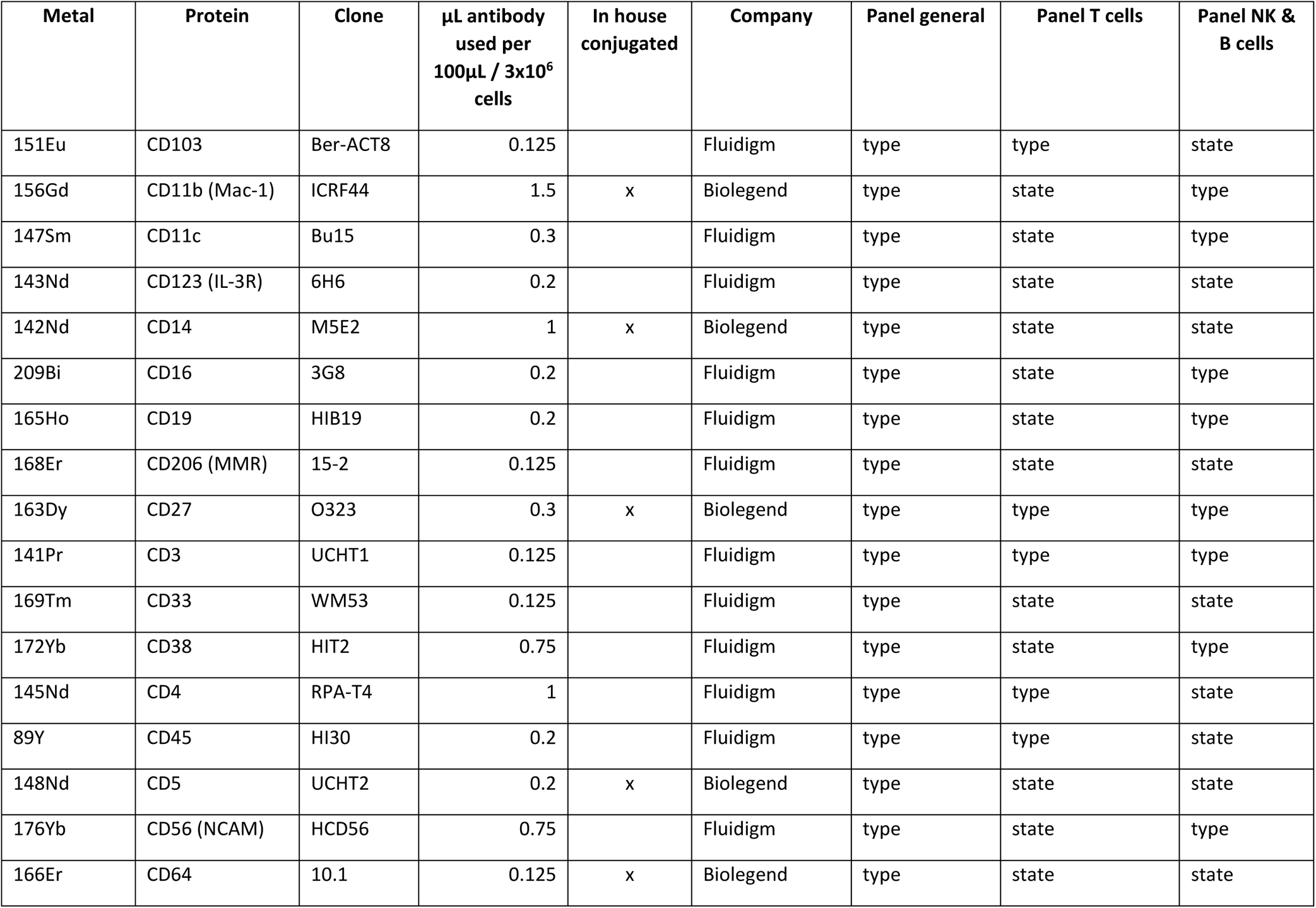
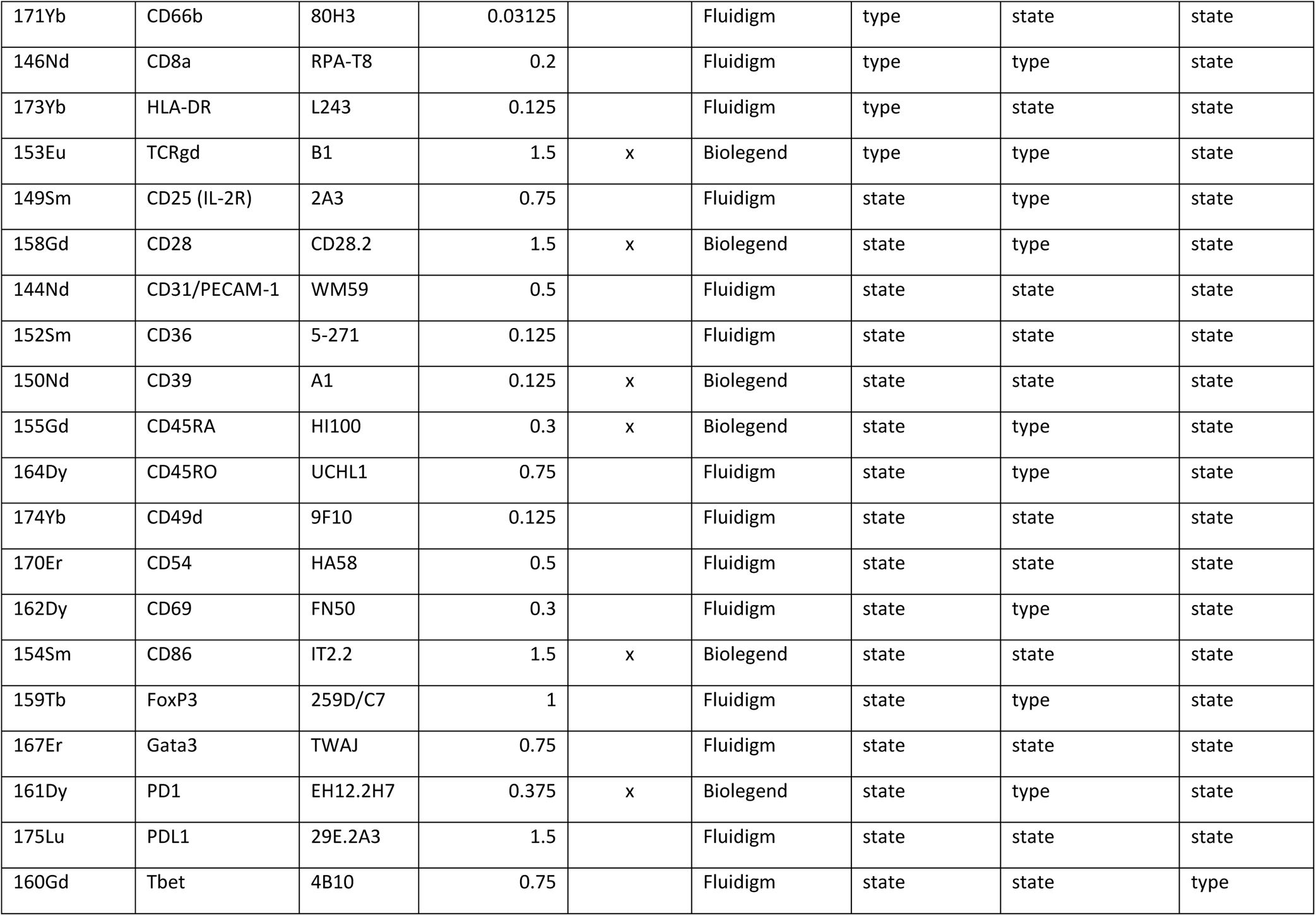
Antibody panel used for CyTOF

**Table S3.** Details of antibodies used for multiplex immunohistochemistry

| Antigen | Species | Dilution | Manufacturer | Cat. Number |
| --- | --- | --- | --- | --- |
| NKp46 (NCR1, CD335) | Rabbit | 1:100 | Abcam | Ab214468 |
| Granzyme K | Rabbit | 1:200 | NovusBio | NBP2-49387 |
| Collagen IV | Rabbit | 1:200 | Abcam | Ab6581 |
| Granzyme B | Rat | 1:100 | Invitrogen | 14-8889-82 |
| CD56 | Rabbit | 1:200 | Abcam | Ab75813 |
| CD45 | Mouse | 1:100 | Dako | M0701 |

## Notes

### Competing Interest Statement

The authors have declared no competing interest.

